# Chromokinesins NOD and KID Use Distinct ATPase Mechanisms and Microtubule Interactions to Perform a Similar Function

**DOI:** 10.1101/520460

**Authors:** Benjamin C. Walker, Wolfram Tempel, Haizhong Zhu, Heewon Park, Jared C. Cochran

## Abstract

Chromokinesins NOD and KID have similar DNA binding domains and functions during cell division, while their motor domain sequences show significant variations. It has been unclear whether these motors have similar structure, chemistry, and microtubule interactions necessary to follow a similar mechanism of force mediation. We used biochemical rate measurements, cosedimentation, and structural analysis to investigate the ATPase mechanisms of the NOD and KID core domains. These experiments and analysis revealed that NOD and KID have different ATPase mechanisms, microtubule interactions, and catalytic domain structures. The ATPase cycles of NOD and KID have different rate limiting steps. The ATPase rate of NOD was robustly stimulated by microtubules albeit its microtubule affinity was weakened in all nucleotide bound states. KID bound microtubules tightly in all nucleotide states and remained associated with the microtubule for more than 100 cycles of ATP hydrolysis before dissociating. The structure of KID was most similar to conventional kinesin (KIF5). Key differences in the microtubule binding region and allosteric communication pathway between KID and NOD are consistent with our biochemical data. Our results support the model that NOD and KID utilize distinct mechanistic pathways to achieve the same function during cell division.

## INTRODUCTION

Kinesins constitute a superfamily of microtubule-based proteins that share a highly conserved catalytic domain.^1, 2^ Kinesin catalytic domains convert energy gained from ATP hydrolysis to a repertoire of mechanical outputs such as stepping along the microtubule lattice or regulating microtubule dynamics.^3^ Small variations in the catalytic domain sequence and structure lead to variations in the ATPase cycles and microtubule interactions, which may explain the large diversity of mechanical functions within the kinesin superfamily.^4^

Chromokinesins NOD (*Drosophila melanogaster*) and KID (human) have convergent mechanical outputs yet their catalytic domain sequences are dissimilar enough to place them in different subfamilies.^5,6^ When taking their C-terminal DNA binding domains and *in vivo* functions into account have they been placed together in the kinesin-10 subfamily.^1, 7^ Given their dissimilar catalytic domain sequence we hypothesize that NOD and KID use divergent ATPase mechanisms to perform their *in vivo* function. NOD and KID, like kinesin-10s in other species, bind chromosome arms during cell division and are responsible for the away from the pole force (polar ejection force) that drives chromosome arm congression at the metaphase plate.^8–13^ How NOD and KID mediate this force is not clear.

Sequence, structure, and kinetic analysis indicate that NOD is monomeric in solution and unlikely to have a stepping mechanism.^14,15^ NOD preferentially binds to microtubule plus-ends^16^, can track microtubule plus-ends *in vivo*^17^, and possesses an unconventional ATPase cycle and microtubule interactions.^14^ In particular, the absence of a conventional neck linker and neck linker binding residues of NOD as well as the unusual weak microtubule interaction in the ATP state speak against a typical stepping mechanism. Additionally, the basal ATPase cycle of NOD is limited by ATP hydrolysis rather than ADP release. Hence the ATPase cycle of NOD is unique and more similar to the non-translocating kinesin MCAK than motile kinesins. This led to the model that NOD acts like a tether between chromosome arms and polymerizing microtubule plus-ends, inhibiting pole-ward fluctuations of chromosome arms (braking mechanism).^14^ Recent *in vivo* and cell extract work showed that NOD not only tracks growing microtubule ends but also moves plus-end directed along the microtubule lattice when artificially dimerized.^17, 18^ NOD may oligomerize upon cargo binding to yield lattice stepping behavior.

Unlike NOD, the catalytic domain and neck linker sequence of KID are more similar to conventional kinesin.^19^ Recently KID has been shown to form homo-oligomeric states in solution^20^ even though other groups failed to show the same.^21, 22^ KID has been shown *in vitro* to be a slow non-processive microtubule plus-end directed motile kinesin.^21–23^ However, given its short distal coiled-coil region, it is yet unclear how a homodimer is capable of forming inter-head tension deemed important for conventional kinesin stepping mechanism.^24, 25^ Little is known about the structure, ATPase cycle, and microtubule interactions of KID.

Whether the microtubule end-binding and braking mechanism or a microtubule lattice stepping mechanism is conserved among the kinesin-10 subfamily to mediate the polar ejection force is not clear. Here we compare the structures of NOD and KID catalytic domains and putative neck linkers, as well as ATPase kinetics and microtubule interactions. We found that the catalytic domain of NOD and KID is markedly different than that of conventional motile kinesins. Under equilibrium conditions NOD binds microtubules unusually weakly, notably in the AMPPNP state, while KID binds unusually tightly, including in the ADP state. Both on and off the microtubule, NOD and KID also follow fundamentally different ATPase cycles. The ATP hydrolysis step limits the basal ATPase rate of NOD which is accelerated upon microtubule binding where ADP release becomes rate limiting. For KID, ADP release limits the basal ATPase cycle. Upon microtubule binding, ADP release is 600-fold stimulated and another yet unknown step becomes rate limiting. We also found that KID is highly chemically processive by remaining associated with the microtubule lattice for over one hundred cycles of ATP hydrolysis before fully dissociating. Our comparison of the catalytic domain crystal structures of NOD and KID predicts differences in how they interact with microtubules, and are consistent with their different kinetic behaviors. Together the data strongly suggest that the catalytic domains of NOD and KID undergo divergent mechanisms to mediate motility or microtubule end tracking that generate the polar ejection force.

## MATERIALS & METHODS

### Experimental conditions

Experiments were performed at 298 K in reaction buffer (20 mM HEPES, pH 7.2 with KOH, 20% (v/v) glycerol, 0.02% (v/v) Tween-20, 2 mM MgCl_2_, 10 mM KCl (low ionic strength) or 100 mM KCl (high ionic strength)) unless otherwise noted. For experiments with polymerized tubulin (microtubules) 20 μM paclitaxel was added to the reaction buffer unless otherwise noted. All concentrations reported are final after mixing.

### Cloning, expression, and purification of KID

We cloned and purified the catalytic domain plus the putative neck linker of human kinesin-like DNA-binding protein (KID; UniProtKB Q14807). The purified protein contained Gly5-Leu379 (MW 41.2 kDa). The coding sequence of the catalytic domain and putative neck linker of human KID was synthesized by polymerase chain reaction (PCR) using human cDNA as a template with primers (5’-TCTAGAGGATCCACGCAGCAGAGGCGACGCG-3’) and (5’-TCTAGAC TCGAGTTACAGGCTCTCATTGGTAAAAGGCCGATTGATC-3’). The synthesized DNA fragment was cloned via BamH*I* and Xho*I* sites into the pJCC04a vector^26^, placing the KID coding sequence immediately downstream of a 6xHis tag and a tobacco etch virus (TEV) protease cleavage site. The 6xHIS-TEV-KID fragment (Nde*I* and Xho*I* digestion) was cloned into the pMALc5x vector (New England Biolabs) via Nde*I* and Sal*I*, placing the maltose binding protein (MBP) coding sequence in frame with the N-terminus of the inserted fragment. The entire coding region was verified by sequencing.

The KID construct was co-expressed with pRILP (Agilent Technologies, Santa Clara, CA) in B834(DE3) cells (EMD Millipore, Darmstadt, Germany). Expression was induced with 0.5 mM isopropyl-β-D-thiogalactopyranoside (IPTG). Cells were lysed in lysis buffer (25 mM NaHPO_4_, pH 7.8, 4 mM MgCl_2_, 0.5 M NaCl, 2 mM DTT, 1 mM EGTA, 2 mM PMSF, 1 mM ATP, 30 mM Imidiazole) with 0.2% (v/v) Triton X-100, 0.05 mg/mL lysozyme, and sonication. Expressed protein was purified by elution from a nickel-nitrilotriacetic acid agarose (Ni-NTA) (Qiagen, Germantown, MD) column with 0.5 M imidazole and buffer exchanged into TEV buffer (25 mM NaHPO_4_, pH 7.8, 4 mM MgCl_2_, 1 mM EDTA, 150 mM NaCl, 2 mM DTT, 0.2 mM ATP, 150 mM sucrose). The MBP solubility tag was cleaved from KID using TEV protease. Cleaved MBP-6xHIS tag was sequestered from KID by Ni-NTA column chromatography. Purified KID was concentrated and buffer exchanged into 20 mM HEPES, pH 7.2 with KOH, 20% (v/v) glycerol, 0.02% (v/v) Tween-20, 4 mM MgCl_2_, 10 mM KCl, 25 μM ATP. Protein concentration was determined by sodium dodecylsulfate polyacrylamide gel electrophoresis (SDS-PAGE) densitometry using bovine serum albumin (BSA) as standards.

### Cloning, expression, and purification of NOD

We cloned and purified the catalytic domain plus twelve C-terminal residues of the *D. melanogaster* no distributive disjunction gene (NOD; UniProtKB P18105). NOD contains the first 330 N-terminal amino acids (Met1-Gln330) followed by Val, Glu, and a 6xHIS tag (MW 37.6 kDa). The coding sequence for the catalytic domain and six adjacent amino acids of NOD was synthesized by PCR using *D. melanogaster* cDNA as a template with primers (5’-TCTAGAATTAAT GGAGGGCGCCAAATTAAGCGC-3’) and (5’-TCTAGAATTAATCTGGCGCGCCACTTGCATCG-3’). The synthesized NOD fragment was digested with Ase*I* and Sal*I* and ligated into the pET24b vector (Novagen, Madison, WI) using the complementary Nde*I* and Xho*I* sites. The entire coding region was verified by sequencing.

NOD was expressed and purified as previously described with few modifications.^27^ Transformed cells were induced with 0.5 mM IPTG at an optical density (A_600_) of ~0.5. Expressed protein was purified from clarified lysate by Ni-NTA column chromatography and was concentrated and buffer exchanged into 20 mM HEPES, pH 7.2 with KOH, 10% (v/v) glycerol, 0.02% (v/v) Tween-20, 2 mM MgCl_2_, 10 mM KCl, 50 μM ATP. Protein concentration was determined by SDS-PAGE densitometry using BSA as standards.

### Preparation of heterodimeric tubulin and stabilized microtubules

Briefly, polymerized tubulin was prepared from bovine brain and stored at 193 K. To obtain depolymerized tubulin, freshly thawed tubulin was incubated on ice 15 min then clarified by centrifugation at 21,130 ×g at 277 K for 15 min. To obtain stabilized microtubules, freshly prepared tubulin was transferred into an equal volume 2× Polymerization Mix (50 mM PIPES pH 6.9, 0.5 mM EGTA, 2 mM GTP, 2.5 mM MgCl_2_, 7.5% DMSO, 50 μM paclitaxel (Acros Organics, Geel, Belgium)) and incubated at 307 K for 45 min. Polymerized and stabilized microtubules were sedimented at 100,000 xg and 298 K for 15 min. Microtubule pellet was suspended in 20 mM HEPES, 20% (v/v) glycerol, 10 mM KCl, 5 mM MgCl_2_ and 50 μM paclitaxel.

To remove the C-terminal tails of α- and β- tubulin, microtubules were treated with 200 μg/mL subtilisin for 45 min at 303 K as described previously.^28^ Proteolysis was stopped with the addition of 5 mM PMSF (Roche Diagnostics). Subtilisin-treated microtubules were spun at 100,000xg for 30 min at 298 K and were resuspended in reaction buffer at 5 mM MgCl_2_, 10 mM KCl, and 50 μM paclitaxel. Tubulin concentration was determined using Coomassie plus Bradford assay reagent (Thermo Fisher Scientific) using BSA as standards.

### Steady-state ATPase assays

Basal, tubulin-, and microtubule-stimulated steady-state ATPase activities were monitored using the NADH coupled assay in reaction buffer.^29–32^ Briefly, for ATP dependent reactions kinesin and tubulin or kinesin and microtubules was diluted to 2× final concentration and mixed with an equal volume of 2x ATP + NADH cocktail (0.4 mM NADH (Acros Organic), 0.5 mM phosphoenolypyruvate (Tokyo Chemical Industry), 5 U/mL rabbit pyruvate kinase (Roche Diagnostics), 8 U/mL lactate dehydrogenase (Sigma-Aldrich)). For tubulin or microtubule dependent reactions kinesin was diluted to 2× final concentration and mixed with an equal volume of 2× tubulin or microtubule, ATP and NADH cocktail. After mixing, reactions were transferred into a 384- well plate in duplicate, pulse centrifuged to 1000 xg, and placed in a microplate spectrophotometer (BioTek) where NADH oxidation was monitored at 340 nm over time. The amount of NADH oxidized (which equals the amount of ADP produced) was calculated via a NADH standard curve (0 μM – 800 μM NADH).

For ATP dependent experiments, the initial velocity (*v*_*o*_) was plotted against the ATP concentration ([ATP]) and the data were fit to the Michaelis-Menten equation (Eq.1), where [E_o_] is the total kinesin concentration, *k*_*cat*_ is the rate of ATP turnover per active site per second, *k*_*m*,ATP_ is the [ATP] at half maximal velocity.

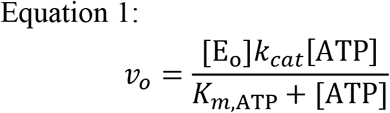

For tubulin or microtubule dependent experiments, the initial velocity (*v*_*o*_) was plotted against the tubulin or microtubule concentration ([T]) and the data were fit to a quadratic equation (Eq.2), where [E_o_] is the total kinesin concentration, k_cat_ is the rate of ATP turnover per active site, *K*_*0.5*_ is the [T] concentration at half maximal velocity, and *k*_*basal*_ is the experimentally determined basal rate of ATP turnover at zero tubulin concentration.

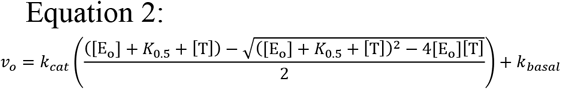

### Cosedimentation assays

Experiments were performed as described with minor modifications.^33^ Briefly, assays were performed at 298 K in reaction buffer and 40-50 μM paclitaxel. To obtain the nucleotide free (apo) state 50 U/mL of alkaline phosphatase (New England Biolabs) was used. To obtain the ADP (Research Products International Corp) and AMPPNP (Sigma Aldrich) states, 1 mM nucleotide was used. Paclitaxel stabilized microtubules were mixed with kinesin and nucleotide and incubated at 298 K for 10-20 min. Complexes of kinesins and microtubules were pelleted using an Optima L-100 XP ultracentrifuge (Beckman Coulter) and a Ti-42.2 fixed-angle rotor at 100,000xg for 10 min. All KID samples and the apo samples for NOD were analyzed using SDS-PAGE densitometry and FIJI software.^34^ The fraction of bound kinesin was plotted against the total microtubule concentration and the data was fit to a quadratic equation (Eq. 3), where *f*_*b,max*_ is the maximal fraction of microtubule bound kinesin, [E_o_] is the total kinesin concentration, and [MT] is the total microtubule concentration.

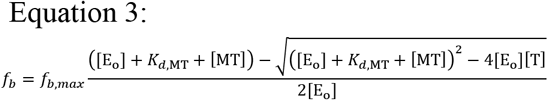

Given the limitations of SDS-PAGE densitometry with NOD, we used the NADH coupled assay to quantify NOD in the supernatant samples. Analyzing three experiments of NOD with AMPPNP using both the SDS-PAGE densitometry and NADH coupled assay methods yielded similar results (Fig. S1). Final results for NOD in the AMPPNP and ADP states reported were analyzed using the NADH coupled assay. Supernatant samples were diluted with reaction buffer and mixed with 5x Master Mix (NADH cocktail, 40 μL paclitaxel, 4 mM ATP, and 2 μM microtubules), transferred to a 384-well plate, pulse centrifuged to 1000xg, and placed in a microplate spectrophotometer (BioTek) where NADH oxidation was monitored at 340 nm over time. The fraction of kinesin bound to microtubules for each condition was calculated (Eq. 4) and was plotted against the microtubule concentration and the data were fit to a quadratic equation (Eq. 3).

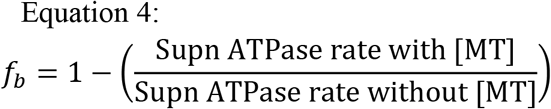

### Stopped-flow experiments

Presteady-state kinetics of mantADP (mant: 2’-(3’)-O-(N– methylanthraniloyl)) release were measured with the SF-300X stopped-flow instrument (KinTek, Snow Shoe, PA). All reactions were performed with 100 mM KCl, 1 μM kinesin, and 1 μM mantADP. The excitation wavelength was set at 356 nm and emitted light was measured through a 400 nm cutoff filter (mant λ_em,max_ = 450 nm). Release of mantADP from the kinesin active site in the presence of tubulin or microtubule was measured by incubating kinesin with mantADP at a 1:1 ratio and rapidly mixing the complex with varying concentrations of tubulin or microtubule plus 1 mM MgATP to chase the tubulin/microtubule•kin•mantADP intermediate. All traces were fit to single exponentials and observed exponential rate of fluorescence decay was plotted as a function of tubulin or microtubule concentration and fit to a hyperbola (Eq. 5), where *k*_*max*_ is the maximum rate of mantADP release, [T] is the total tubulin or microtubule concentration, *k*_*d*,mADP_ is the [T] at half maximal velocity (½ *k*_*max*_), and *k*_*basal*_ is rate of mantADP release at zero tubulin or microtubule.

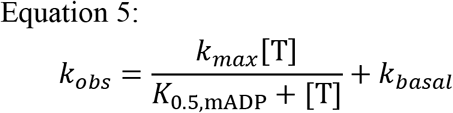

### Malachite green assay

To measure mantATPase activity, a malachite green assay was used as described previously with minor modifications.^35^ Reactions were performed in reaction buffer and 2 mM MgCl_2_ at 1 mM mantATP and 10 mM KCl except for microtubule dependent reactions where 0.25 mM mantATP and 100 mM KCl were used. For basal mantATPase activity, NOD was diluted to a final concentration of 1 μM and KID diluted to 2 μM in reaction buffer. For microtubule-stimulated activity, NOD and KID were diluted to a final concentration of 0.1 μM and for tubulin-stimulated activity NOD was diluted to 1.5 μM. For microtubule dependent reactions NOD was diluted to 0.3 μM. Tubulin and microtubule-stimulated reactions were performed with either 1 μM microtubules and 20 μM paclitaxel or 2 μM tubulin. To initiate the reaction 15 μL of 2 mM mantATP was mixed with an equal volume of 2x kinesin at room temperature. After increasing times of incubation (up to 12 min) the entire reaction volume was quenched by rapidly mixing with 7.5 μL of 5 N HCl. After exactly 5 min post quenching 450 μL of malachite green solution was added and incubated at room temperature. During incubation 150 μL of each reaction was transferred to a 96-well plate in triplicate. At exactly 12.5 min post mixing the absorbance at 650 nm was obtained with a microplate spectrophotometer (BioTek). The reported absorbance values were converted into phosphate concentrations using a phosphate standard curve (12.5 μM – 330 μM NaPO_4_, pH 7.2).

### Cloning, expression, purification, and crystallization of KID

We cloned and purified the catalytic domain plus the putative neck linker of human KID (Pro40-Lys400) for crystallization. The coding sequence of KID was cloned into the pET28-MHL vector as 6His-TEV-KID. The KID construct was expressed in BL21-CodonPlus cells and induced with 0.5 mM IPTG. Cells were lysed in binding buffer (1X PBS, 0.5 M NaCl, 5 mM Imidazole, 5 mM BME, pH 7.0) with 0.5% (v/v) protease inhibitor cocktail (Sigma), 1600 units Benzonase (Sigma), and a microfluidizer. Expressed protein was purified first by elution from Ni-NTA beads with elution buffer (20 mM Hepes, 0.5 M NaCl, 300mM Imidazole, pH 7.0, 5 mM BME, 2 mM MgCl_2_) and then by size exclusion chromatography (Superdex 75, GE-Healthcare) in size exclusion buffer (20 mM Hepes, pH 7.0, 500 mM NaCl, 1 mM TCEP). KID was 95% pure as judged by SDS-PAGE and concentrated to 32 mg/ml in size-exclusion buffer. Concentrated KID was mixed with 5 molar fold of MgADP, 1% (w/w) Endoproteinase Glu-C V8 protease for crystallization. Well-diffracting crystals formed at 291 K within 1–2 days when using 3.2 M NaCl, 0.1 M Tris pH 8.5 as crystallization buffer. Crystals were cryo protected with the addition of ethylene glycol in reservoir solution and flash frozen with liquid nitrogen.

**Figure 1.**
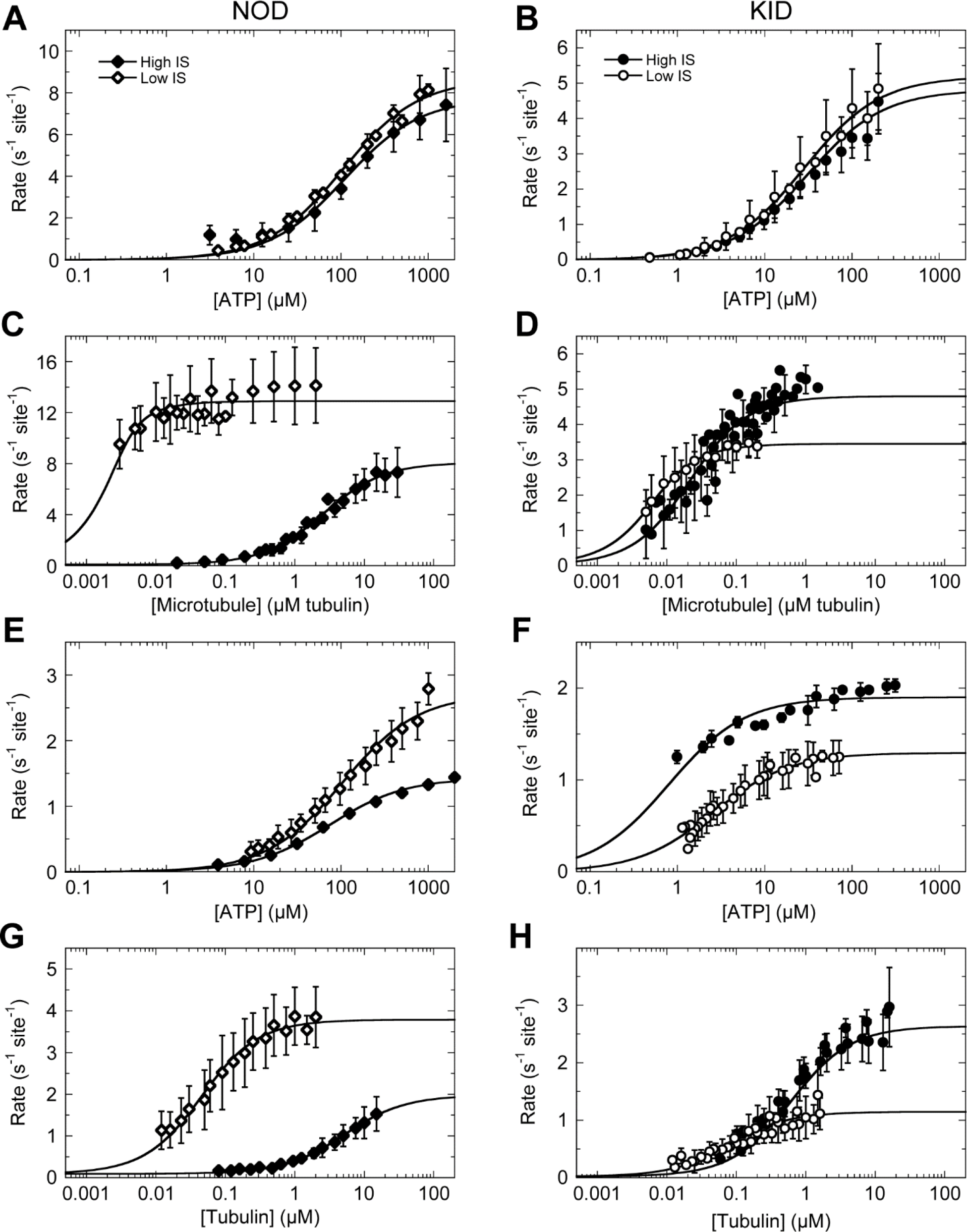
Microtubule and Tubulin-Stimulated Steady-state ATPase of NOD and KID. Error bars represent the standard deviation across independent experiments (n = 3-14). Reported error for each constant represents the standard error obtained from non-linear least squares analysis. A. ATP dependent microtubule-stimulated ATPase of NOD. At low ionic strength (♢): 2 μM microtubules; 43 nM NOD; *k*_*cat*_ = 8.7 ± 0.2 s-1; *K*_*m*,ATP_ = 105 ± 7 μM. At high ionic strength (♦): 20 μM microtubules; 30 nM NOD; *k*_*cat*_ = 7.8 ± 0.4 s-1; *K*_*m*,ATP_ = 110 ± 20 μM. B. ATP dependent microtubule-stimulated ATPase of KID. At low ionic strength (○): 2 μM microtubules; 60 nM KID; *k*_*cat*_ = 5.2 ± 0.2 s-1; *K*_*m*,ATP_ = 28.3 ± 2.7 μM. At high ionic strength (●): 2 μM microtubules; 60 nM KID; *k*_*cat*_ = 4.8 ± 0.1 s-1; *K*_*m*,ATP_ = 30.8 ± 2.5 μM. C. Microtubule dependent ATPase of NOD. At low ionic strength (♢): 2 mM ATP; 3 nM NOD; *k*_*cat*_ = 12.8 ± 0.3 s-1; *K*_*0.5*,MT_ = 0.0005 ± 0.0002 μM. At high ionic strength (♦): 2 mM ATP; 20 nM NOD; *k*_*cat*_ = 8.0 ± 0.2 s-1; *k*_*0.5,MT*_ = 2.6 ± 0.2 μM. D. Microtubule dependent ATPase of KID. At low ionic strength (○): 1 mM ATP; 5 nM KID; *k*_*cat*_ = 3.43 ± 0.05 s-1; *K*_*0.5*,MT_ = 0.0034 ± 0.0003 μM. At high ionic strength (●): 1 mM ATP; 5 nM KID; *k*_*cat*_ = 4.8 ± 0.1 s-1; *K*_*0.5*,MT_ = 0.019 ± 0.002 μM. E. ATP dependent tubulin stimluated ATPase of NOD. At low ionic strength (♢): 1 μM tubulin; 10 nM NOD; *k*_*cat*_ = 2.7 ± 0.1 s-1; *K*_*m*,ATP_= 103 ± 12 μM. At high ionic strength (♦): 15 μM tubulin; 100 nM NOD; *k*_*cat*_ = 1.43 ± 0.02 s-1; *K*_*m*,ATP_ = 74 ± 5 μM. F. ATP dependent tubulin-stimulated ATPase of KID. At low ionic strength (○): 1 μM tubulin; 40 nM KID; *k*_*cat*_ = 1.30 ± 0.02 s-1; *K*_*m*,ATP_ = 2.6 ± 0.1 μM. At high ionic strength (●): 15 μM tubulin; 30 nM KID; *k*_*cat*_ = 1.90 ± 0.04 s-1; *K*_*m*,ATP_ = 0.8 ± 0.1 μM. G. Tubulin dependent ATPase of NOD. At low ionic strength (♢): 2 mM ATP; 10 nM NOD; *k*_*cat*_ = 3.69 ± 0.07 s-1; *K*_*0.5*,TUB_= 0.040 ± 0.004 μM. At high ionic strength (♦): 3 mM ATP; 60 nM NOD; *k*_*cat*_ = 1.88 ± 0.06 s-1; *K*_*0.5*,TUB_ = 5.2 ± 0.4; μM. H. Tubulin dependent ATPase of KID. At low ionic strength (○): 1 mM ATP; 10 nM KID; *k*_*cat*_ = 1.12 ± 0.05 s-1; *K*_*0.5*,TUB_ = 0.08 ± 0.01 μM. At high ionic strength (●): 1 mM ATP; 10 nM KID; *k*_*cat*_ = 2.6 ± 0.1 s-1; *K*_*0.5*,TUB_ = 0.61 ± 0.08 μM.

### Data collection and structure refinement

Diffraction data were collected on a rotating copper anode x-ray source and reduced with the HKL suite.^36^ The structure was solved by molecular replacement with the program PHASER^37^ and coordinates from PDB entry 3B6U. The programs DM^38^ and RESOLVE^39^ were used for phase modification. SCWRL^40^ was used for the adjustment of model sidechains to conform with the target amino acid sequence. COOT^41^ was used for interactive model rebuilding. REFMAC^42^, PHENIX^43^ and BUSTER^44^ were used for restrained model refinement. PyMOL^45^ was used for model visualization and preparation of figures. X-ray data collection and refinement statistics are given in Table S1. The PDB ID code is 6NJE.

**Figure 2.**
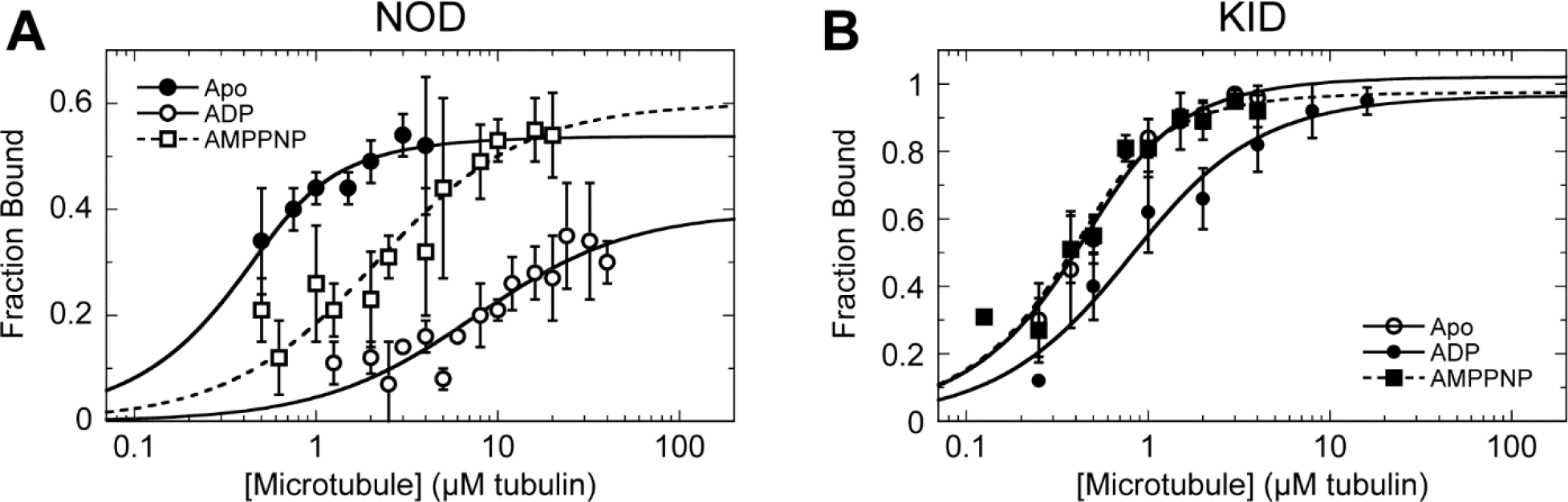
Cosedimentation of NOD• microtubule and KID• microtubule under Different Nucleotide States. Error bars represent the standard deviation across independent experiments (n = 3-8). Reported error for each constant represents the standard error obtained from non-linear least squares analysis. A. The fraction of NOD bound in the microtubule pellet was plotted against the microtubule concentration. For NOD+Alk Phos (◊) the *K*_*d*,MT(apo)_ NOD+AMPPNP (◻) the *K*_*d*,MT(AMPPNP)_ = 1.9 ± 0.5 μM; *f*_*b,max*(AMPPNP)_ = 0.6. For NOD+ADP (○)the *K*_*d*,MT(ADP)_ = 7.3 ± 2.0 μM; *f*_*b,max*(ADP)_ = 0.4. B. The fraction of KID bound in the microtubule pellet was plotted against the microtubule concentration. For KID+Alk Phos (◊) the *K*_*d*,MT(apo)_ = 0.16 ± 0.04 μM; *f*_*b,max*(apo)_ =1.0. For KID+AMPPNP (◻) the *K*_*d*,MT(AMPPNP)_ = 0.11 ± 0.04 μM; *f*_*b,max*(AMPPNP)_ = 1.0. For KID+ADP (○) the *K*_*d*,MT(ADP)_ = 0.5 ± 0.1 μM; *f*_*b,max*(ADP)_ = 1.0.

### Homology modeling of NOD

Homology models of NOD were created using SWISS-MODEL.^46^ The NOD catalytic domain and neck linker were modeled using the full length NOD sequence as querry and known kinesin structures with docked neck linkers from a large variety of subfamilies as templates [Kinesin-1: KIF5C (PDB: 3X2T), KIF5B (PDB: 1MKJ); Kinesin-2: KIF3B (PDB: 3B6U), KIF3C (PDB: 3B6V); Kinesin-3: KIF1A (PDB:1I6I), KIF13B (PDB:5ZBR); Kinesin-4: KIF4 (PDB: 3ZFC); Kinesin-5: Eg5 (PDB: 3HQD); Kinesin-6: MKLP2 (PDB: 5ND4); Kinesin-7: CENP-E (PDB: 1T5C); Kinesin-8: KIF18A (PDB: 5OCU); Kinesin-12: KIF15 (PDB: 4BN2)]. When multiple structures were available we selected structures with larger portions of their neck sequence ordered (when applicable) or at random. We only considered structures from motile kinesins with an N-terminal motor domain.

## RESULTS

### Expression and purification of kinesin motors

We cloned, expressed, and purified the catalytic domain and putative neck linker of human KID and of *D. melanogaster* NOD (Fig. S2). KID was expressed with a maltose binding protein solubility tag and a 6x histidine purification tag, which were removed via TEV protease cleavage during purification. NOD was purified with a C-terminal 6× histidine tag. The expression and purification of NOD and KID yielded approximately 14.7 mg and 0.66 mg per liter of media respectively, with >95% purity as determined by SDS-PAGE densitometry (Fig. S2).

### The NOD-microtubule/tubulin interactions are unusually sensitive to ionic strength

We measured the ATP, microtubule, and tubulin dimer dependent steady-state ATPase activity at low and high ionic strengths (Fig. 1, Table 1). The activity of each kinesin is robustly stimulated by microtubules and tubulin.

**Figure 3.**
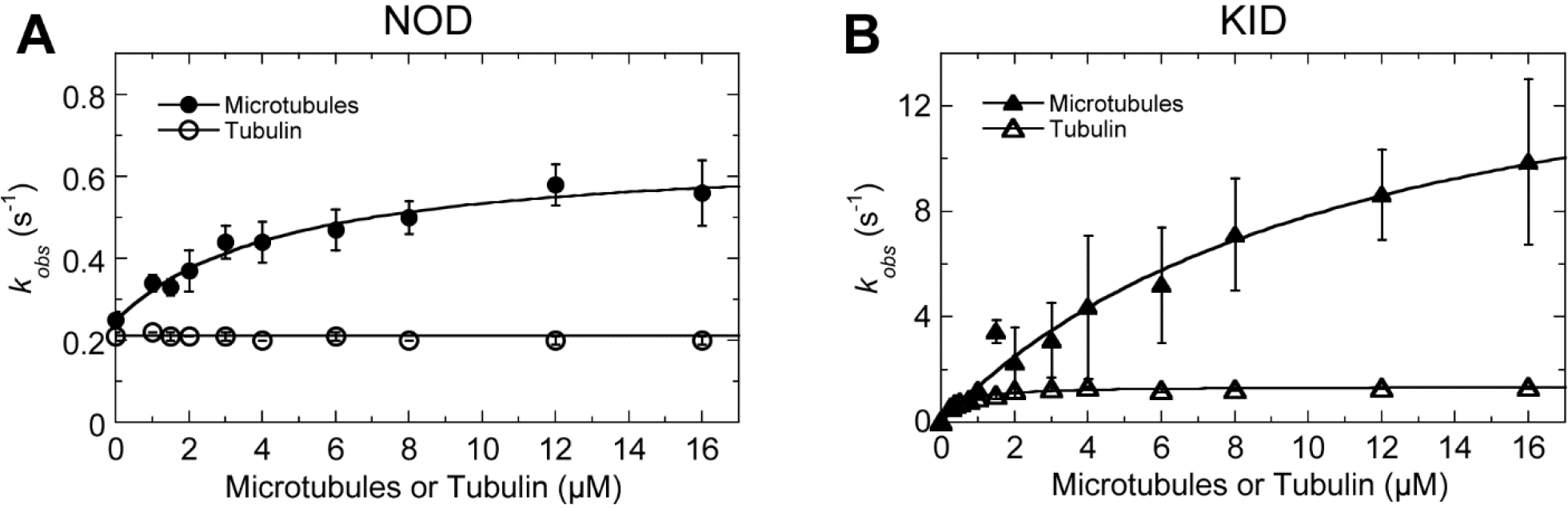
mantADP release kinetics of NOD and KID. All experiments were performed in reaction buffer at high ionic strength. Error bars represent the standard deviation across independent experiments. Reported error for each constant represents the standard error obtained from non-linear least squares analysis. A) Microtubule (▲) and tubulin (△) dependent mantADP release kinetics for NOD: Basal = 0.24 ± 0.03 s^−1^; microtubule-stimulated = 0.66 ± 0.03 s^−1^; *K*_*0.5*,MT_ = 4.6 ± 0.9 μM; Tubulin-stimulated = 0.21 ± 0.01 s^−1^. B) Microtubule (▲) and tubulin (△) dependent mantADP release kinetics for KID: Basal = 0.028 ± 0.005 s^−1^; microtubule-stimulated = 16.9 ± 2.5 s^−1^; *K*_*0.5*,MT_ = 11.6 ± 3.1 μM; Tubulin-stimulated = 1.34 ± 0.03 s^−1^; *K*_*0.5*,TUB_ = 0.43 ± 0.04 μ

Additionally, both kinesins have physiologically relevant affinities for ATP in the presence of microtubule and tubulin that are minimally affected by ionic strength. Significant differences between NOD and KID emerge at their microtubule lattice and tubulin interactions.

While the affinity of KID for microtubules decreased only 8-fold with a 10-fold increase in ionic strength, the microtubule affinity for NOD decreased 5,200-fold. Similarly, the affinity for tubulin of KID decreased only 6-fold while the affinity of NOD decreased 128-fold. This strong salt dependence of the microtubule/tubulin and NOD interaction predicts that polymer binding of NOD is much more dependent on polar interactions compared to KID and other kinesins.^47,48^

The weak microtubule and tubulin affinities of NOD at high ionic strength (*K*_*0.5*,MT_ = 2.6 ± 0.3 μM; *K*_*0.5*,TUB_ = 5.2 ± 0.3 μM) indicate that the ATPase cycles of NOD in the presence of microtubules or tubulin are dominated by weakly interacting intermediate states such as the NOD•ADP state. By comparison the tight microtubule and tubulin interactions of KID at high ionic strength (*K*_*0.5*,MT_ = 0.019 ± 0.018 μM; *K*_*0.5*,TUB_ = 0.6 ± 0.1 μM) indicate that tightly bound states dominate its ATPase cycle.

**Table 1:**
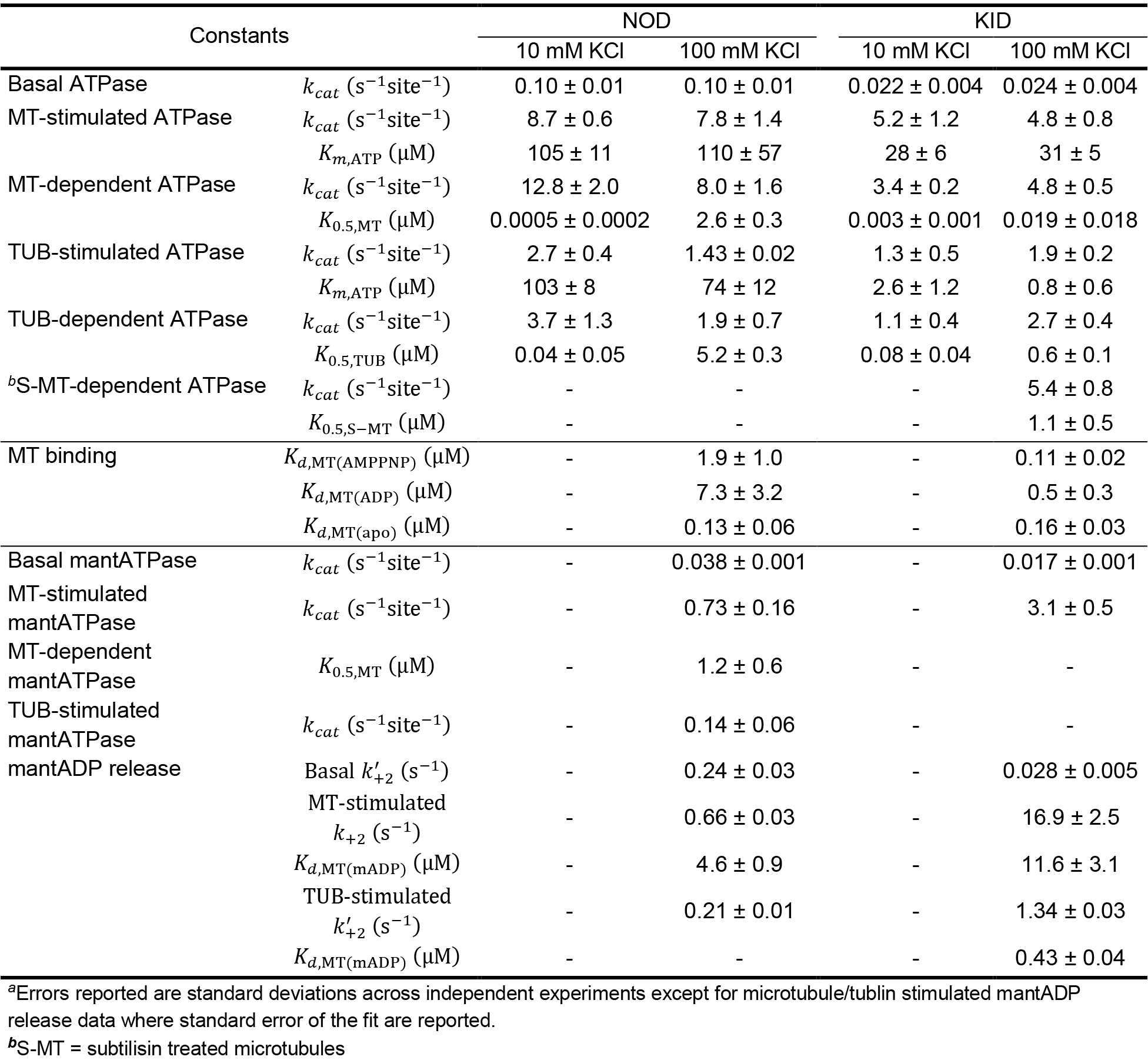
Comparison of Kinetic Constants for NOD and KID^*a*^

| Constants |  | NOD |  | KID |  |
| --- | --- | --- | --- | --- | --- |
|  |  | 10 mM KCl | 100 mM KCl | 10 mM KCl | 100 mM KCl |
| Basal ATPase | $k_{cat}$ ($s^{-1}$site$^{-1}$) | $0.10 \pm 0.01$ | $0.10 \pm 0.01$ | $0.022 \pm 0.004$ | $0.024 \pm 0.004$ |
| MT-stimulated ATPase | $k_{cat}$ ($s^{-1}$site$^{-1}$) | $8.7 \pm 0.6$ | $7.8 \pm 1.4$ | $5.2 \pm 1.2$ | $4.8 \pm 0.8$ |
| | $K_{m,ATP}$ (μM) | $105 \pm 11$ | $110 \pm 57$ | $28 \pm 6$ | $31 \pm 5$ |
| MT-dependent ATPase | $k_{cat}$ ($s^{-1}$site$^{-1}$) | $12.8 \pm 2.0$ | $8.0 \pm 1.6$ | $3.4 \pm 0.2$ | $4.8 \pm 0.5$ |
| | $K_{0.5,MT}$ (μM) | $0.0005 \pm 0.0002$ | $2.6 \pm 0.3$ | $0.003 \pm 0.001$ | $0.019 \pm 0.018$ |
| TUB-stimulated ATPase | $k_{cat}$ ($s^{-1}$site$^{-1}$) | $2.7 \pm 0.4$ | $1.43 \pm 0.02$ | $1.3 \pm 0.5$ | $1.9 \pm 0.2$ |
| | $K_{m,ATP}$ (μM) | $103 \pm 8$ | $74 \pm 12$ | $2.6 \pm 1.2$ | $0.8 \pm 0.6$ |
| TUB-dependent ATPase | $k_{cat}$ ($s^{-1}$site$^{-1}$) | $3.7 \pm 1.3$ | $1.9 \pm 0.7$ | $1.1 \pm 0.4$ | $2.7 \pm 0.4$ |
| | $K_{0.5,TUB}$ (μM) | $0.04 \pm 0.05$ | $5.2 \pm 0.3$ | $0.08 \pm 0.04$ | $0.6 \pm 0.1$ |
| <sup>b</sup>S-MT-dependent ATPase | $k_{cat}$ ($s^{-1}$site$^{-1}$) | - | - | - | $5.4 \pm 0.8$ |
| | $K_{0.5,S-MT}$ (μM) | - | - | - | $1.1 \pm 0.5$ |
| MT binding | $K_{d,MT(AMPPNP)}$ (μM) | - | $1.9 \pm 1.0$ | - | $0.11 \pm 0.02$ |
| | $K_{d,MT(ADP)}$ (μM) | - | $7.3 \pm 3.2$ | - | $0.5 \pm 0.3$ |
| | $K_{d,MT(apo)}$ (μM) | - | $0.13 \pm 0.06$ | - | $0.16 \pm 0.03$ |
| Basal mantATPase | $k_{cat}$ ($s^{-1}$site$^{-1}$) | - | $0.038 \pm 0.001$ | - | $0.017 \pm 0.001$ |
| MT-stimulated mantATPase | $k_{cat}$ ($s^{-1}$site$^{-1}$) | - | $0.73 \pm 0.16$ | - | $3.1 \pm 0.5$ |
| MT-dependent mantATPase | $K_{0.5,MT}$ (μM) | - | $1.2 \pm 0.6$ | - | - |
| TUB-stimulated mantATPase | $k_{cat}$ ($s^{-1}$site$^{-1}$) | - | $0.14 \pm 0.06$ | - | - |
| mantADP release | Basal $k'_{+2}$ ($s^{-1}$) | - | $0.24 \pm 0.03$ | - | $0.028 \pm 0.005$ |
| | MT-stimulated $k_{+2}$ ($s^{-1}$) | - | $0.66 \pm 0.03$ | - | $16.9 \pm 2.5$ |
| | $K_{d,MT(mADP)}$ (μM) | - | $4.6 \pm 0.9$ | - | $11.6 \pm 3.1$ |
| | TUB-stimulated $k'_{+2}$ ($s^{-1}$) | - | $0.21 \pm 0.01$ | - | $1.34 \pm 0.03$ |
| | $K_{d,MT(mADP)}$ (μM) | - | - | - | $0.43 \pm 0.04$ |

<sup>a</sup>Errors reported are standard deviations across independent experiments except for microtubule/tubulin stimulated mantADP release data where standard error of the fit are reported.
<sup>b</sup>S-MT = subtilisin treated microtubules

### NOD and KID show different modulation of microtubule affinity under various nucleotide conditions

To investigate the microtubule lattice interaction of NOD and KID in different nucleotide states, we performed kinesin•microtubule cosedimentation assays under MgADP, MgAMPPNP, and no nucleotide (apo) conditions (Fig. 2, Table 1). To obtain nucleotide-free conditions, we treated each kinesin with alkaline phosphatase to cleave ADP to adenosine and inorganic phosphate (P_i_). Fraction bound was calculated from the unbound fraction in the supernatant by SDS-PAGE analysis or NADH-coupled ATPase activity. NOD binds microtubules tightly at high ionic strength in the apo state (*K*_*d*,MT_ = 0.13 ± 0.06 μM), and more weakly in the AMPPNP (*K*_*d*,MT_ = 1.9 ± 1.0 μM) and ADP states (*K*_*d*,MT_ = 7.3 ± 3.2 μM). The relatively weak microtubule affinity of NOD in the AMPPNP state is consistent with previous results albeit our results show a tighter binding in the AMPPNP state compared to the ADP state.^27, 49^ In comparison KID binds tightly in all nucleotide states with the ADP state being weaker than the apo and AMPPNP states (*K*_*d*,MT(ADP)_ = 0.5 ± 0.1 μM, *K*_*d*,MT(apo)_ = 0.16 ± 0.04 μM, *K*_*d*,MT(AMPPNP)_ = 0.11 ± 0.04 μM).

Interestingly only a fraction of NOD will bind excess microtubules (*f*_*b,max*(ADP)_ = 0.4 ± 0.1; *f*_*b,max*(apo)_ = 0.5 ± 0.1; *f*_*b,max*(AMPPNP)_ = 0.6 ± 0.1). Contaminating unpolymerized tubulin does not account for the unbound fraction (data not shown). Partial binding has been observed previously with KIF5A and Kar3 motor domains and likely is a motor specific ionic strength effect.^48, 50^ In comparison the entire KID population binds excess microtubules (*f*_*b,max*_ = 1.0 ± 0.1).

### MantADP release of NOD is weakly stimulated by microtubules compared to KID

MantADP release kinetics was measured by rapidly mixing kinesin•mantADP with increasing concentrations of microtubules or tubulin plus excess ATP in a stopped flow instrument. The basal mantADP release rate of NOD (0.24 ± 0.03 s^−1^) was only 3-fold accelerated by microtubules (0.66 ± 0.03 s^−1^) and not at all by tubulin (0.21 ± 0.01 s^−1^) (Fig. 3, Table 1). By comparison the basal mantADP release rate of KID (0.028 ± 0.005 s^−1^) was 600-fold accelerated by microtubules (16.9 ± 2.5 s^−1^) and 50-fold by tubulin (1.34 ± 0.03 s^−1^).

### MantATPase rates are slowed for NOD but not for KID

To ensure that mant-labelled nucleotide kinetics are representative of unlabeled nucleotide, we measured mant-ATPase steady-state kinetics using the malachite green assay. The basal mantATPase kinetics of NOD is 3-fold slower compared to its basal ATPase rate while its microtubule-, and tubulin-stimulated kinetics were 10-fold slower (Fig. 1, Table 1, Fig. S3). The affinity for microtubules of NOD in the presence of mantATP was 2-fold tighter than in the presence of non-labelled ATP (Fig 1, Table 1, Fig. S4). The mantATPase kinetics of KID, on the other hand, were consistent with its ATPase kinetics suggesting that its mant-nucleotide kinetics were representative of kinetics with unlabeled nucleotide.

### KID was highly chemically processive at the microtubule lattice

The *K*_*d*,MT(ADP)_ for KID as observed in our equilibrium experiments was tight at 0.5 μM. Under steady-state conditions the rate for ATP turnover at 1 μM microtubule is ~5 s^−1^ for KID. However, in our pre-steady state ADP release experiments the *K*_*d*,MT(mADP)_ was 11.6 μM and the rate of ADP release at 1 μM microtubule was well under 1 s^−1^. We interpret this data as KID initially weakly interacting with microtubules. Once KID is bound to microtubules it remains associated with the microtubule for many rounds of ATP hydrolysis. Further support comes from the unusually high chemical processivity of the motor head of KID (Table 2). With a *K*_*bi*_ ratio of 139 the KID catalytic domain hydrolyses over one hundred ATPs per microtubule encounter. We hypothesized that KID relies heavily on tubulin’s C-terminal tails (CTTs) to remain associated with the microtubule without fully dissociating. We treated microtubules with subtilisin to cleave off the CTTs of microtubules and observed that the affinity of KID for subtilisin treated microtubules decreased 46-fold from *K*_*0.5*,MT_ = 0.024 ± 0.018 μM to *K*_*0.5*,S-MT_ = 1.1 ± 0.5 μM (Fig. S5, Table 1). Interestingly, the new *K*_*bi,S-kcat*_ is 4.9, much more in line with other monomeric kinesins and, assuming the *K*_*bi,mADP*_ is not significantly affected by CTTs, the new *K*_*bi*_ ratio now is 3.3.

**Table 2:** *K*_*bi*_ Values of Monomeric Kinesin Constructs

| Constants | Kinesin-1 | Kinesin-5 | Kinesin-10 |  | Kinesin-14 |  |
| --- | --- | --- | --- | --- | --- | --- |
|  | *DmKHC*<sup>51</sup> | *HsEg5*<sup>52</sup> | *HsKID* | <sup>a</sup>*HsKID*<sup>TUB</sup> | *DmNCD*<sup>53</sup> | *ScKar3*<sup>54</sup> |
| $K_{bi,kcat}$ | 6.8 | 8 | 208 | 5.4 | 0.22 | 0.08 |
| $K_{bi,mADP}$ | 2.1 | 10 | 1.5 | 1.5 | 0.13 | 0.07 |
| $K_{bi,kcat}/K_{bi,mADP}$ | 3.2 | 0.8 | 139 | 3.6 | 1.7 | 1.1 |

<sup>a</sup>Values derived from experiments with tubulin instead of microtubules.

**Figure 4.**
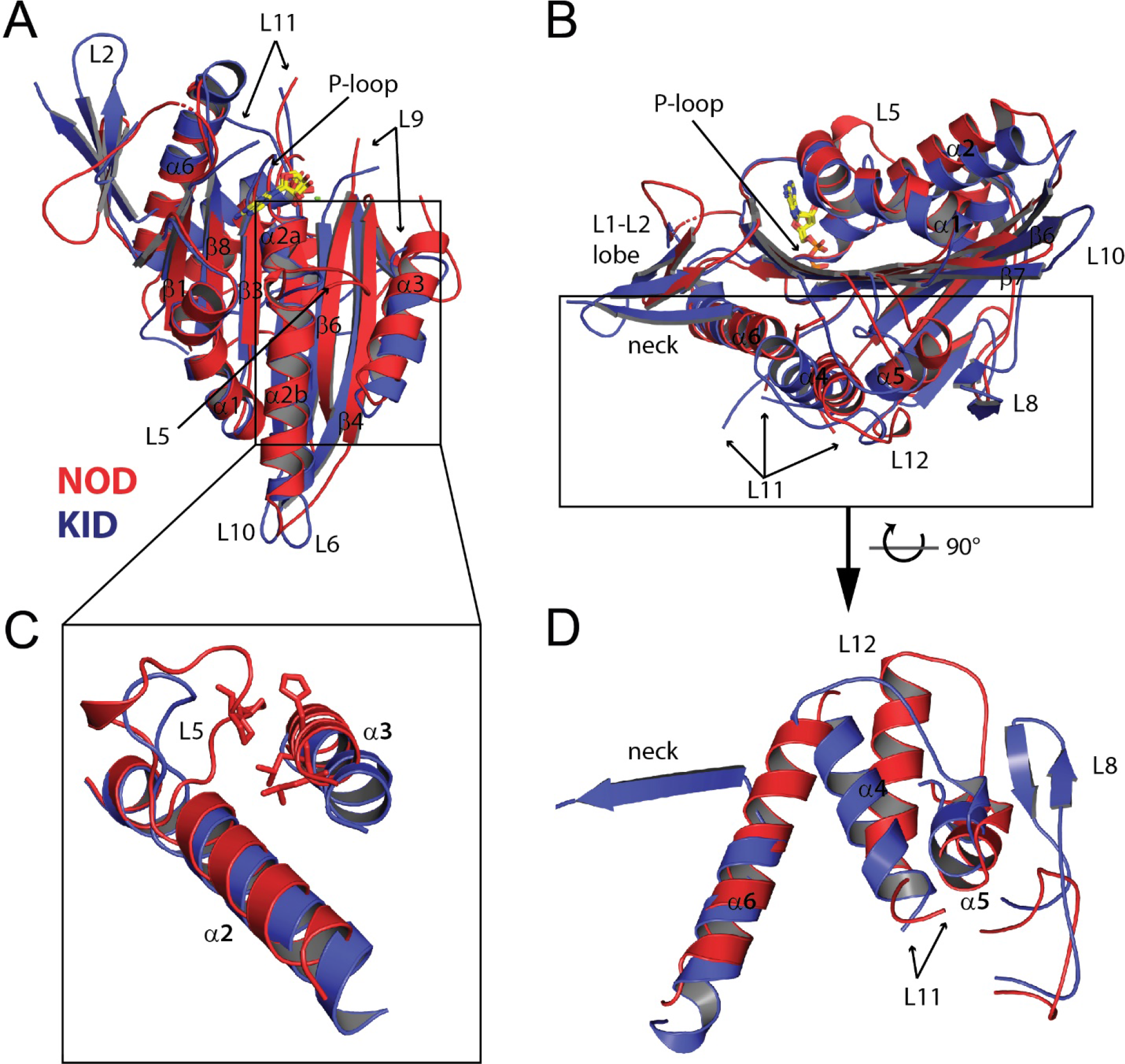
Structural Model Comparison of NOD and KID. Superposition of KID•ADP and NOD•ADP via the P-loop. A) Top view. B) Side view. C) The extended L5 interaction with α3 helix of NOD compared to KID. D) Comparison of the microtubule binding interface of NOD and KID.

The high chemical processivity was also absent in the presence of tubulin. The *K*_*0.5*,TUB_ for steady state ATPase was 0.5 ± 0.1 μM, which was comparable to the *K*_*0.5*,TUB_ for pre-steady-state ADP release (0.4 ± 0.2 μM). The chemical processivity of KID in the presence of tubulin is 3.6, which was like other monomeric kinesins further supporting that the high chemical processivity of KID in the presence of microtubule was lattice specific (Table 2). The chemical processivity for NOD when using mant-nucleotide kinetics was slightly higher when compared to other monomeric kinesins (7 vs ≤ 3) but significantly lower than that of KID, further highlighting significant differences in microtubule interaction of NOD and KID during their ATPase cycles.

### Structural comparison between KID and NOD

We solved the X-ray crystal structure of the KID catalytic domain in the ADP state (Fig. 4A, Table S1). We compared the X-ray crystal structure of the KID•ADP catalytic domain structure with that of NOD•ADP (PDB entry 3DC4).^27^ The two structures were aligned using the P-loop with a root mean square deviation (RMSD) of 1.6 Å over 236 Cα positions (PDBe Fold v2.59). The largest differences between the KID and NOD structures were observed in the microtubule binding regions of L8 and helix α4, switch-1 containing loop L9, the half of the β-sheet near L8 (β4, β5, β6, and β7), β1-L1, L5, L10, and helix α6 (Fig. 4A-4B).

**Figure 5.**
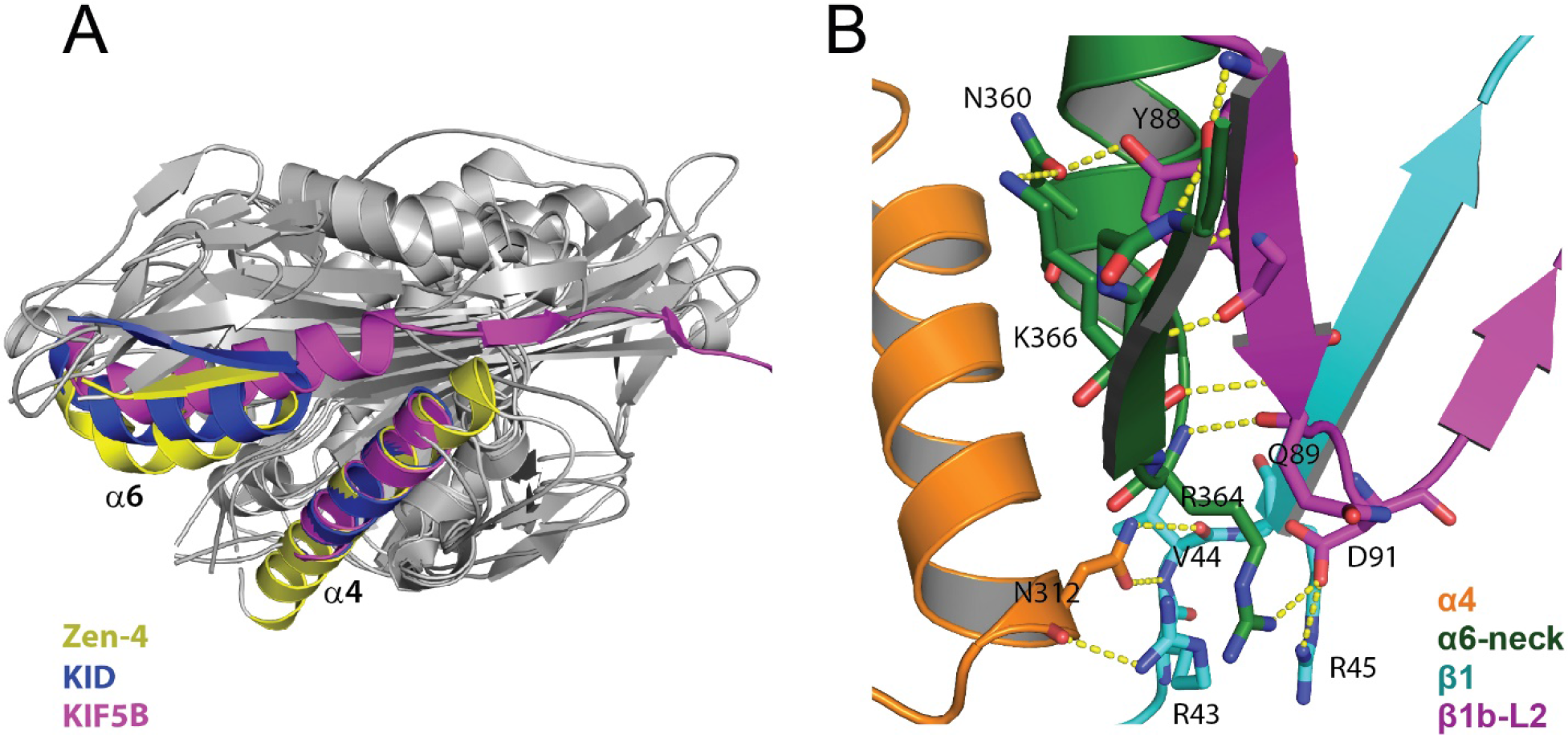
Neck Linker Orientation of KID. A) Superposition of KID, Zen4 (5X3E), and KIF5B (1MKJ) via the helix α4 to highlight α6-neck orientation relative to helix-α4. B) Polar interactions between α4, α6-neck, β1, and the L1/L2 lobe that stabilize the backward orientation of the neck linker of KID.

A notable difference was evident in L5 at the top of the catalytic domain (Fig. 4C). The L5 of NOD, but not KID, extends and interacts with α3. Striking differences between KID and NOD exist in the microtubule-binding L8 and helix α4 (Fig. 4D). Typically α4 shows two dominant conformations: the “ADP-like” (up = C-terminal end of α4 away from microtubule) and the “ATP-like” (down = C-terminal end of α4 toward the microtubule as well as counterclockwise rotation of ~12 Å toward α6) conformations (Fig. S6A).^55^The α4 of KID is in the ADP-like “up” conformation whereas the α4 of NOD has a unique conformation similar only the kinesin-13 KIF2C as described previously (Fig. S6B).^27^

While the crystal structure construct of NOD did not include the putative neck linker residues its α4 helix is oriented away from helix α6, avoiding steric clashes and allowing α6 to fully extend (Fig. 4D). However, the α4 helix in the NOD•ADP structure is still too close to the core (“up” position) to allow putative neck linker docking. NOD does not have highly conserved neck linker residues found in motile kinesins originally thought to underlie its non-motility.^19, 49^ However, NOD does have key conserved residues at the α6-neck junction necessary for neck linker docking as well as conserved residues that make up the hydrophobic pocket (Fig. S7).^56^ To evaluate whether neck linker docking is possible we created homology models of the catalytic domain of NOD with its putative neck linker residues using known kinesin structures from ten different kinesin subfamilies with docked neck linkers (Fig. S7). In all cases the L320 of NOD docks into the hydrophobic pocket (neck linker cleft), suggesting that although NOD has a non-conventional extended neck-linker sequence it may retain the ability to dock and contribute to force generation along the microtubule lattice.

The α6 helix and neck linker orientation of KID vary from known ADP-like kinesin structures. The α6 helix is partially melted with the neck linker docked back to form a β-sheet with β1a/β1b in the L1/L2 lobe (Fig. 4D, Fig. 5). The α6/neck structure of KID is very similar to the recently published nucleotide-free structure of Zen-4 (PDB entry 5X3E), suggesting that this conformation is likely a general feature of kinesins (Fig. 5A).^57^ Guan *et al.* published data suggesting that this feature is particularly prominent in the nucleotide free leading head of dimeric kinesins when tension is pulling the neck linker backwards.^57^ In KID, the N-terminal junction of β1 and the L1/L2 lobe come together to interact with the C-termini of α4 and α6 to stabilize the partially melted α6 and backward conformation of the neck linker (Fig. 5B). Residue N312 in α4 makes three contacts at the β1 junction (R43, V44). R364 in the melted α6 makes a sharp turn and forms a bond network with β1b/L2 (Q89, D91), and β1 (R45). The backbone of residues S365 – I369 now form β-sheet interactions with β1b and the side chain of K366 interacts with N360 of α6 and the side chain of Y88 of β1b.

## DISCUSSION

We compared the catalytic domain of NOD and KID and their putative neck linker residues by analyzing their ATPase kinetics, microtubule interactions, and crystal structures. We found that the catalytic domains of NOD and KID markedly differ from conventional motile kinesins. NOD binds microtubules unusually weakly in the AMPPNP state while KID binds unusually tightly in the ADP state. NOD and KID also follow fundamentally different ATPase cycles on and off the microtubule with different rate limiting states. We also found that KID is highly chemically processive. Our comparison of the catalytic domain crystal structures of NOD and KID predicts differences in how they interact with microtubules and are consistent with their different kinetic behaviors.

### ADP release is rate limiting for the NOD•microtubule but not the KID•microtubule ATPase mechanism

The basal mantADP release rate of NOD (0.24 ± 0.03 s^−1^) was fast compared to its basal mantATPase rate (0.038 ± 0.001 s^−1^) (Fig. 3, Table 1). Additionally, tubulin did not stimulate mantADP release of NOD yet was able to activate its mantATPase rate. Together these results show that mantADP release was not limiting the basal mantATPase cycle of NOD as previously seen.^27^ By comparison the basal mantADP release rate of KID (0.028 ± 0.005 s^−1^) was in good agreement with its basal mantATPase (0.017 ± 0.001 s^−1^) and ATPase (0.024 ± 0.004 s^−1^) rates indicating that ADP release limits the basal ATPase cycle of KID.

The microtubule-stimulated mantADP release rate for NOD (0.66 ± 0.03 s^−1^) corresponds well with its microtubule-stimulated mantATPase rate (0.73 ± 0.16 s^−1^), suggesting that mantADP release was limiting the microtubule-stimulated mantATPase cycle of NOD (Fig. 3, Table 1). In contrast, the microtubule-stimulated mantADP release rate of KID (16.9 ± 2.5 s^−1^) was fast compared to its microtubule-stimulated mantATPase and ATPase rates (3.1 ± 0.5 s^−1^; 4.6 ± 0.5 s^−1^) indicating that ADP release does not limit the microtubule-stimulated ATPase cycle of KID. Together our results show that NOD and KID follow fundamentally different ATPase cycles on and off the microtubule.

The tubulin-stimulated mantADP release rate of NOD (0.21 ± 0.01 s^−1^) approaches the rate of its tubulin-stimulated mantATPase rate (0.14 ± 0.06 s^−1^) suggesting that ADP release was at least partially rate limiting (Fig. 3, Table 1). Similarly, the tubulin-stimulated mantADP release rate of KID (1.34 ± 0.03 s^−1^) was in good agreement with its tubulin-stimulated ATPase rate (1.9 ± 0.2 s^−1^) indicating that ADP release rate also limits the tubulin-stimulated ATPase cycle of KID.

### The rate limiting step of NOD was accelerated by microtubules

The NOD construct used in this study differs from our previously published construct (NOD318).^27^ The main feature of the current construct was tighter microtubule binding. While both constructs bound tightly in the apo state (≤ 0.1 μM), NOD318 showed significantly weaker binding in the presence of nucleotides. Under steady-state ATP turnover conditions, NOD318 bound tenfold weaker under low ionic strength compared to the current construct (Fig. S8). The reduced binding affinity of NOD318 can explain the lack of microtubule-stimulated ATP hydrolysis previously published.^27^ However, values derived from mantADP release experiments are comparable between the two constructs leading us to believe that the rate limiting step has not changed. Given that the rate limiting step of NOD (ATP hydrolysis) was stimulated by microtubules, it occurs while NOD was microtubule bound.

The difference in microtubule affinity may largely be due to the negatively charged N-terminal FLAG tag of NOD318 that may interfere with binding to the negatively charged microtubule lattice. For the new construct, we exchanged the N-terminal FLAG tag with a C-terminal 6x histidine tag and extended the sequence by 12 amino acids into the neck region that aligned with the neck region of the KID construct that contained conserved neck linker residues.

### NOD and KID have unusual nucleotide specific microtubule binding states

The ability of kinesin to step along the microtubule lattice depends on cyclical tight and weak interaction states that are determined by the active site nucleotide state. Typically, motile kinesins bind to microtubule tightly in the apo and the ATP-like states and weakly in the ADP state.^58, 59^ The catalytic domains of NOD and KID diverge from this general rule.

Weakly bound states dominate the ATPase cycle of NOD as suggested by the observed high *K*_*0.5*,MT_ at high ionic strength (Fig. 1). This was consistent with ADP release limiting the microtubule-stimulated ATPase cycle and unusually weak microtubule affinity in the ATP-like AMPPNP state (Fig. 2). Our observation was consistent with the literature albeit the AMPPNP state was still more tightly bound to microtubules than the ADP state (2 ± 1 μM vs 7 ± 3 μM).^27,49^ The only other kinesin with evidence of a weak ATP state is the non-motile kinesin MCAK.^60, 61^ This behavior is inconsistent with a conventional stepping mechanism along the microtubule lattice, and NOD failed to produce *in vitro* motility in our hands (data not shown) and in the hands of others.^18, 49^ Interestingly, longer constructs of NOD do have plus-end directed motility *in vitro* but only in the presence of cell extract or when artificially dimerized and purified from insect cells.^18^ We suspect that the ATP state of NOD binds microtubules sufficiently tight for stepping either when induced via cellular factors and/or because dimerization and gating of the heads induce a tightly microtubule bound state.

Unlike NOD, KID binds microtubules tightly in the ADP state as well as the AMPPNP and apo states (Fig. 2). It is therefore not surprising that KID binds microtubules tightly during its ATPase cycle (Fig. 1). Similar behavior has been observed for a monomeric construct of the kinesin-3, KIF1A.^62^ Monomeric KIF1A binds tightly in all nucleotide states but was able to partially dissociate from its tight binding site and diffuse along the microtubule lattice.^63^ Dimerization triggers a change in motor behavior that allows KIF1A to processively walk along the microtubule lattice.^64^ KID can form a weak oligomeric state in solution with its short distal coiled-coil domain.^20^ Like KIF1A, KID may require dimerization to obtain motility competent microtubule binding behavior.^23^

### The KID catalytic domain has a high chemical processivity

Microtubule affinity of KID•ADP under equilibrium conditions was tight (*K*_*d*,MT(ADP)_ = 0.5 ± 0.3 μM) and in the presence of 1 μM microtubules the rate of steady-state ATP turnover was ~5 s^−1^ (Fig. 1, Fig. 2). Surprisingly, the microtubule affinity of KID as observed in the pre-steady state ADP release experiments was weak (*K*_*0.5*,MT_ = 11.6 μM) and in the presence of 1 μM microtubules the rate of ADP release was <1 s^−1^ (Fig. 3). We interpret this paradox as KID initially weakly interacting with microtubules in the ADP state. Once KID is bound to the microtubule it remains associated for many rounds of ATP hydrolysis (high chemical processivity).

Consistent with our interpretation, we calculated an unusually high chemical processivity of the motor head of KID (Table 2). With a *K*_*bi,MT*_ ratio of 139 the KID catalytic domain hydrolyses over one hundred ATPs per microtubule encounter. We hypothesized that KID relies on tubulin’s C-terminal tails (CTTs) to remain associated with the microtubule without fully dissociating. Associating with CTTs allows KID to bind weakly to its binding site in the ADP state without fully dissociating. Consistent with this idea the microtubule affinity decreased nearly 50-fold when we removed the CTTs with subtilisin treatment (*K*_*0.5*,MT_ = 0.024 ± 0.018 μM to *K*_*0.5*,S-MT_ = 1.1 ± 0.5 μM) (Fig. S4, Table 1). The high chemical processivity of KID was microtubule specific as its chemical processivity in the presence of tubulin was low and similar to other monomeric kinesin constructs (Table 2).

### NOD and KID differ in their microtubule binding domains

The P-loop aligned structures of KID and NOD show significant differences in their microtubule binding motifs (Fig. 4B,D). The structure of KID was most similar to kinesin-1 ADP-like structures with the α4 helix oriented in the “up” position, occluding the neck linker binding pocket and coinciding with a shortening of the α6 helix and a backward docking of the neck linker (Fig. 5). The α6 helix of NOD was extended and oriented away from its α4 helix by ~32° more than KID. However, the α4 helix of NOD was not far enough in a “down” position to allow neck linker docking (Fig. 5B). Homology modeling of NOD with its unconventional neck linker showed that neck linker docking into the hydrophobic pocket was possible (Fig. S7).

Kinesin structures can be divided into three subdomains analogous to myosins: the lower subdomain (L11-α4-L12-α5) and the upper-and N-terminal subdomains which are derived by dividing the central β-sheet at the beta strands anchoring the P-loop (β3) and the switch-2 loop (β7).^65^ Upon docking KID and NOD structures onto the microtubule lattice (aligning helix α4 of NOD, KID and kinesin-1 bound to the microtubule), the N-terminal and upper subdomain of NOD are rotated clockwise compared to KID by ~25.5° (Fig. S9A). NOD rotated while avoiding clashes with the microtubule lattice due to its missing hairpin in L8 (Fig. S9B). NOD and KID, like other ADP-like structures, have a shortened α4 helix and an open polymer cleft. In both structures, clashes between the lower subdomain and the microtubule lattice are observed (α6 of NOD also clashes with the lattice) (Fig. S9C). These putative clashes may underlie the weak polymer affinity of ADP-like structures.^65^ KID and NOD are therefore predicted to bind microtubules weakly in the ADP state. We observed weak binding for NOD•ADP in our equilibrium experiments (Fig. 2). This prediction supports our model derived from our ADP release data and chemical processivity observations that KID•ADP interacts weakly with the microtubule lattice but then remains associated via the negatively charged CCTs of tubulin.

### NOD and KID have key structural differences in their allosteric communication pathways

Differences in the active site, especially the switch motifs/loops, raise the possibility of differences in allosteric communication. The nucleotide position and binding, phosphate interactions, and magnesium coordination are well conserved between KID and NOD. The respective switch-2 motifs in L11 are similarly positioned at the active site in the “open” position (Fig. S10). However the remainders of the switch-2 loop adopt distinct conformations. Whereas the switch-1 motifs in L9 are in the open conformation for both kinesins, their conformations diverge and interact differently with the switch-2 motifs (Fig. S10).

The L5 of NOD in the N-terminal subdomain extends and interacts with α3 in the upper subdomain (Fig. 4C), similar to Eg5 inhibition by monastrol.^27^ Monastrol binds at the interface between L5 and α3 and inhibits ADP release, leading to slow kinesin•microtubule complex formation. Consistent with this observation, NOD exhibits unusually limited stimulation of ADP release upon microtubule binding and a weak microtubule affinity in the ADP state (Table 1). The L5-α3 interaction in NOD represents an atypical contact between the N-terminal and upper subdomains and may therefore interfere with the rearrangement of the subdomains upon the engagement of the linchpin residue by the microtubule. Hence this interaction interferes with the allosteric communication pathway between the polymer binding site and the active site and therefore was thought to affect ATPase activity and motility.^27, 66^ By comparison the L5 of KID was too short to interact with its α3. Accordingly, KID shows strongly accelerated ADP release and tight microtubule affinity (Table 1). Together these structural observations support the divergent ATPase mechanisms and microtubule interactions of NOD and KID, as observed in our biochemical assays.

### Kinetic and structural data are consistent with published in vivo findings

The catalytic domain of NOD alone did not bind microtubules *in vivo*, but remained diffuse in the cytosol.^18^ Longer constructs containing a short low probability coiled-coil domain weakly interacted with microtubules but did not enrich at the plus ends. Only artificial dimerization showed robust microtubule interaction, motility and microtubule plus-end tracking. Our *in vitro* data corroborate weak microtubule binding of the catalytic domain and a biochemistry inconsistent with motility. It is likely that NOD oligomerization was induced by cargo binding, leading to sufficiently tight microtubule binding for motility and end-tracking. This could be an effective strategy to ensure that NOD interacts with a microtubule lattice/ends only after it was bound to chromosomes.

KID has multiple functions during cell division. In prometaphase, KID is mainly associated along the spindle until metaphase when it relocates mainly to the chromosomes.^10, 67^ This switch correlates with two distinct functions, its role in spindle stability and morphogenesis and its role in the polar ejection force and chromosome arm congression. For spindle stability, KID binds tightly to spindle microtubules, which does not require its ATPase and DNA binding activities.^67, 68^ Phosphorylation and other cellular factors lead to a weakening of microtubule binding and loading onto chromosomes where KID produces the PEF.^13, 67, 69–71^ Our observed tightly bound catalytic domain with high chemical processivity may reflect its biochemistry during spindle formation. Only upon cargo binding can KID efficiently dimerize and become motile.

## ASSOCIATED CONTENT

Supplemental Figures and Tables (separate file).

### Accession numbers

UniProtKB Accession ID: KID Q14807; NOD P18105. RCSB

Accession Code: KID 6NJE

### Author Contributions

B.C.W. and J.C.C. designed the study, performed the research, and wrote the manuscript. H.P. initiated and supervised KID structure determination. H.Z. oversaw expression, purification and crystallization of KID. W.T. prepared an earlier version of the KID atomic model.

B.C.W. refined the KID crystal structure model and performed the structure analysis.

### Funding

This work was supported by Indiana University and the National Science Foundation Grant MCB 1614514 (to J.C.C.). The authors declare that they have no conflicts of interest with the contents of this article. The content is solely the responsibility of the authors and does not necessarily represent the official views of the National Institute of Health. The SGC is a registered charity (number 1097737) that receives funds from AbbVie, Bayer Pharma AG, Boehringer Ingelheim, Canada Foundation for Innovation, Eshelman Institute for Innovation, Genome Canada through Ontario Genomics Institute [OGI-055], Innovative Medicines Initiative (EU/EFPIA) [ULTRA-DD grant no. 115766], Janssen, Merck KGaA, Darmstadt, Germany, MSD, Novartis Pharma AG, Ontario Ministry of Research, Innovation and Science (MRIS), Pfizer, São Paulo Research Foundation-FAPESP, Takeda, and Wellcome.

## Acknowledgements

We thank Jeff Ewer for purifying the NOD protein and Caleb Starr for initial characterizations of the KID protein. We thank Farrell MacKenzie for cloning and Yang Shen for purifying KID protein for crystallization. We thank Claire Walczak and Stephanie Ems-McClung for valuable discussion.

## Abbreviation

MT: microtubule
TUB: free heterodimeric tubulin
MBP: maltose binding protein
TEV: tobacco etch virus
IPTG: ispropyl-β-D-thiogalactopyranoside
BME: 2- mercaptoethanol
Ni-NTA: nickel-nitrilotriacetic acid agarose
6xHIStag: 6x histidine purification tag
PCR: polymerase chain reaction
SDS-PAGE: sodium dodecyl sulfate polyacrylamide gel electrophoresis
BSA: bovine serum albumin
PEF: polar ejection force
CTTs: C-terminal tails

