## Supplemental Info for "Chromokinesins NOD and KID Use Distinct ATPase Mechanisms and Microtubule Interactions to Perform a Similar Function"

### Supplemental Material

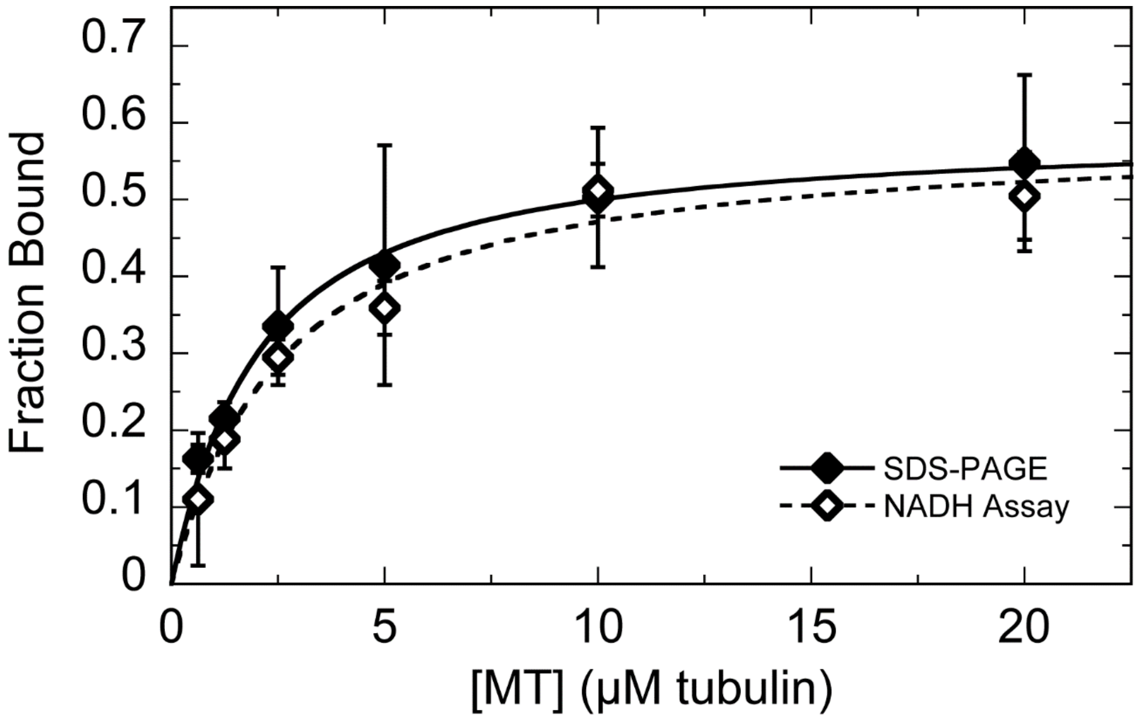


**Figure S1. Comparison of NADH-assay and SDS-PAGE analysis of NOD•microtubule cosedimentation assays.** Cosedimentation assay in the presence of AMPPNP was performed as described. Error bars represent the standard deviation across three independent experiments. Fraction bound was obtained by analysis of supernatants by SDS-PAGE (♦) and NADH-coupled ATPase assay (◊). For SDS-PAGE analysis: $K_{d,\mathrm{MT(AMPPNP)}}$ = 1.7 ± 0.3 μM (error of fit = 0.2); $f_{b,max}$(AMPPNP) = 0.6. For NADH-coupled assay: $K_{d,\mathrm{MT(AMPPNP)}}$ = 2.3 ± 0.7 (error of fit = 0.5); $f_{b,max}$(AMPPNP) = 0.6. Errors are standard deviations.

Table S1: Summary of Crystallographic Data Refinement Statistics.

| Refinement | |
|---|---|
| Resolution (Å) | 35.5 – 2.2 |
| $R_{cryst}$ / $R_{free}^{a}$ (%) | 20.0 / 25.8 |
| No. Atoms (protein/solvent) | 2333 / 90 |
| R.M.S. Deviations | |
| Bond Lengths (Å) | 0.007 |
| Bond Angle (°) | 0.950 |
| Avg. *B* Factors (Å$^2$) | 39.4 |
| Ramachandran – Procheck | |
| [A,B,L] | 285 (98.3%) |
| [a,b,l,p] | 5 (1.7 %) |
| [~a,~b,~l,~p] | 0 (0.0 %) |

[a]$R_{free}$ is for 5% of total reflections not included in the refinement.

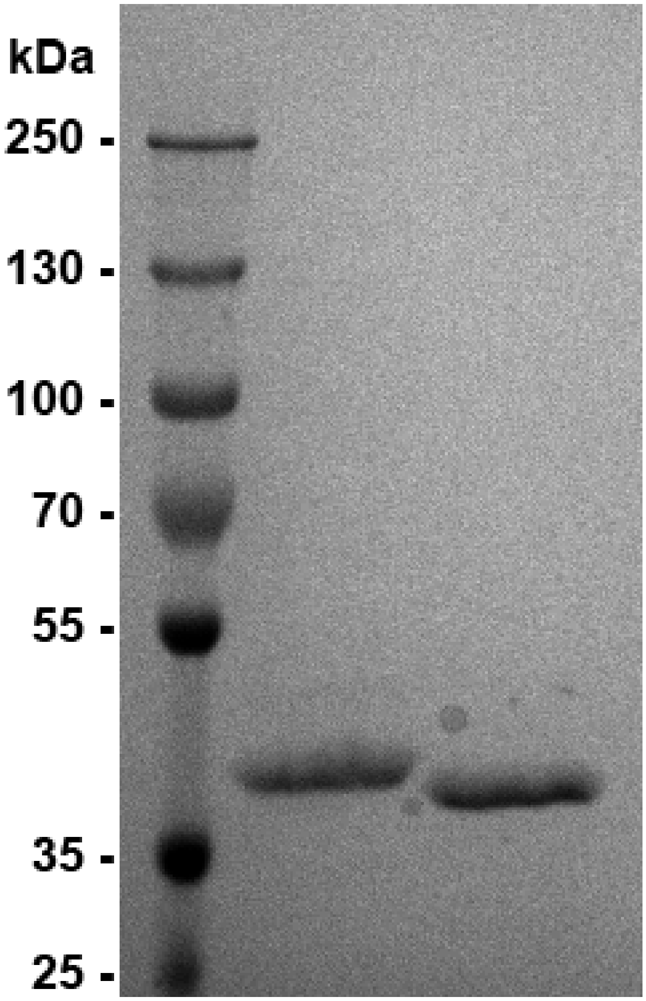


**Figure S1. SDS-PAGE analysis of purified KID and NOD.** From left to right: ladder, 52 pmol KID, 52 pmol NOD.

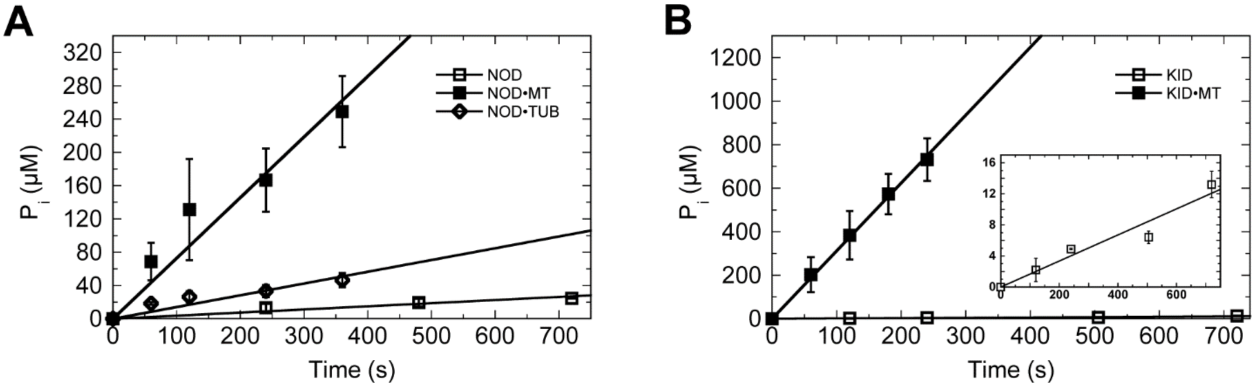


**Figure S3. mantATPase kinetics of NOD and KID.** All experiments were performed in reaction buffer at low ionic strength. Error bars represent the standard deviation across independent experiments. Reported error for each constant represents the standard error obtained from non-linear least squares analysis. A. mantATPase for NOD: Basal (□; $k_{cat}$ = 0.038 ± 0.003 $s^{-1}$), microtubule-stimulated (■; $k_{cat}$ = 0.73 ± 0.06 $s^{-1}$), and tubulin-stimulated (◊; $k_{cat}$ = 0.14 ± 0.02 $s^{-1}$). B. mantATPase for KID: Basal (■; $k_{cat}$ = 0.017 ± 0.001 $s^{-1}$; n=3; inset) and microtubule-stimulated (□; $k_{cat}$ = 3.12 ± 0.04 $s^{-1}$; n=3).

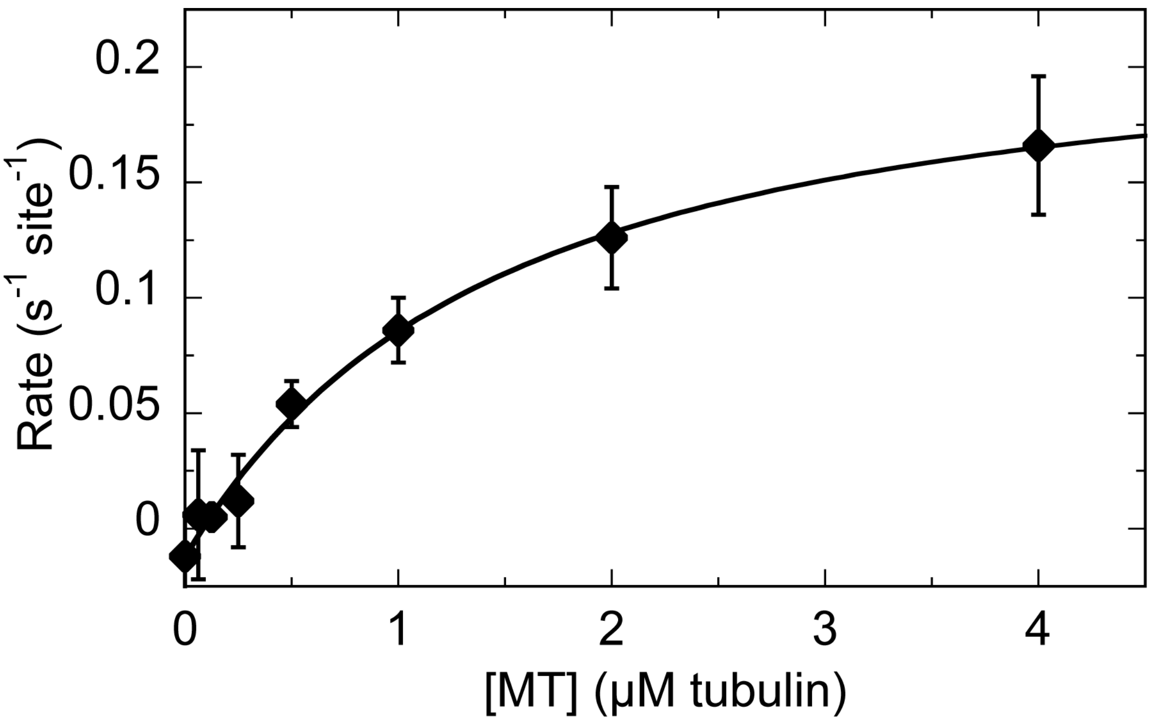


**Figure S4. Microtubule dependent mantATPase of NOD.** Error bars represent the standard deviation across independent experiments (n=3). Reported error for each constant represents the standard error obtained from non-linear least squares analysis. Experiments were performed in reaction buffer at high ionic strength, 0.25 mM mantATP, 0.3 µM NOD, 2 mM $MgCl_2$, and 20 µM paclitaxel. $K_{0.5,\mathrm{MT}} = 1.2 \pm 0.2$ µM.

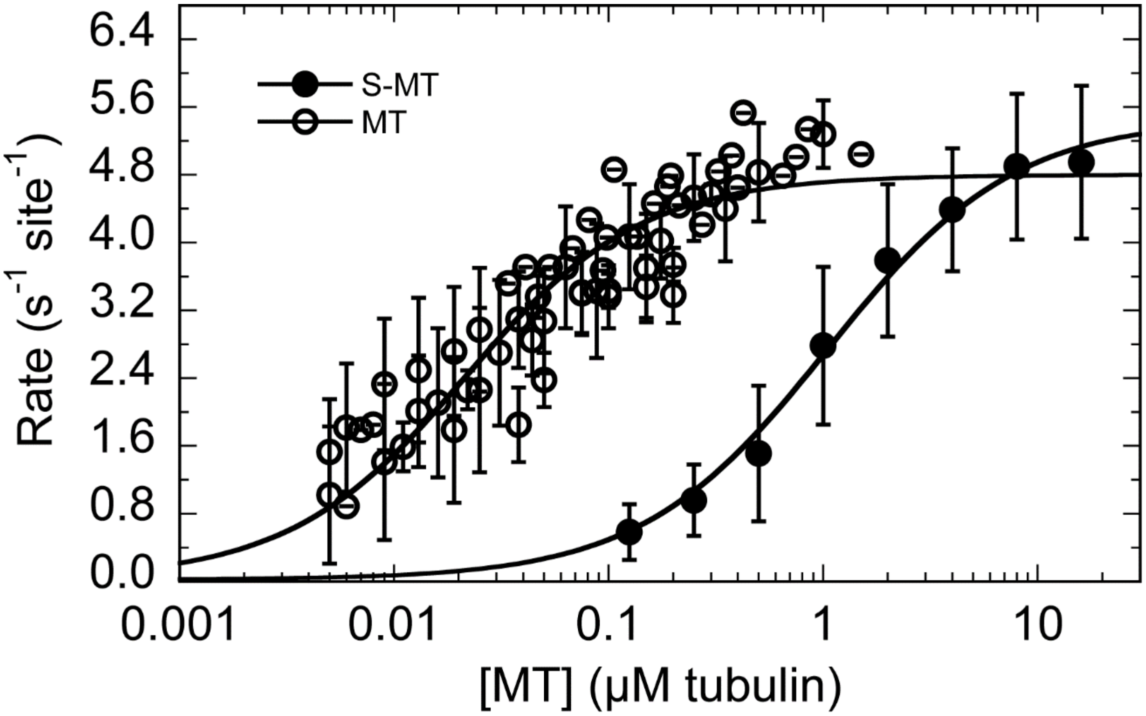


**Figure S5. Subtilisin treated microtubule dependent steady-state ATPase of KID.** Untreated microtubule (MT) data (see Figure 1D) is included as reference. Experiments were performed in reaction buffer at high ionic strength, 1 mM ATP, 0.03 µM KID, 2 mM $MgCl_2$, and 20 µM paclitaxel. Error bars represent the standard deviation across three independent experiments. $k_{cat}$ = 5.4 ± 0.8 $s^{-1}$ (error of fit: 0.2); $K_{0.5,\text{S-MT}}$ = 1.1 ± 0.5 µM (error of fit: 0.1). Reported errors are standard deviations.

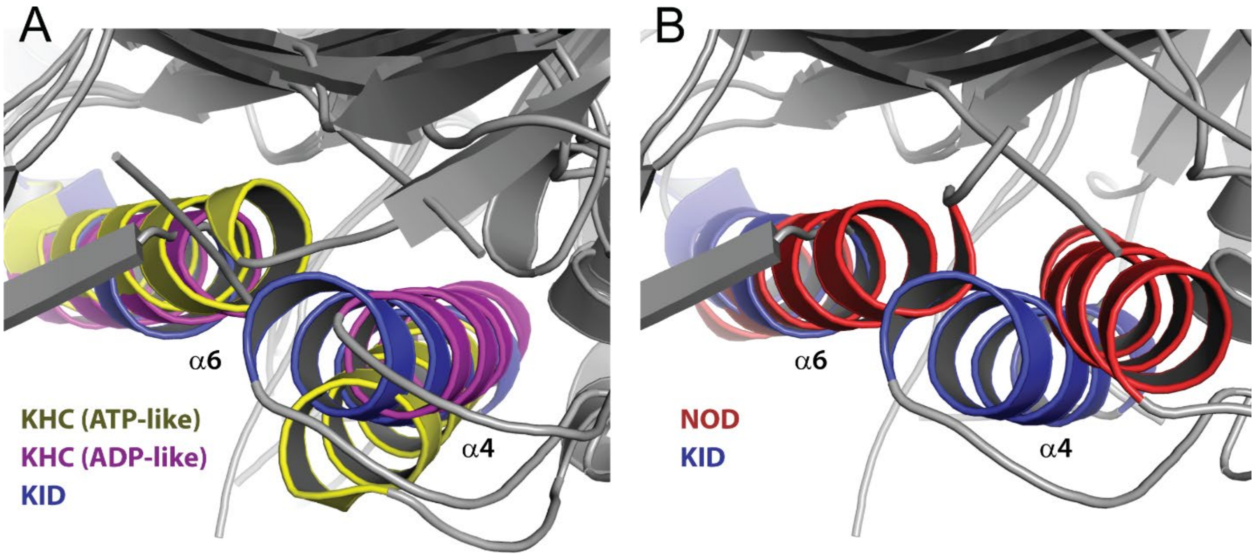


**Figure S6. Helix-α4 orientation of NOD and KID.** A) The α4 of KID (blue) is in the ADP-like position when compared to KHC (kinesin-1 ADP-like (1BG2) and ATP-like (1MKJ)). B) The α4 of NOD has a unique orientation pointing away from α6 and is in between the ADP-like and ATP-like position relative to the core.

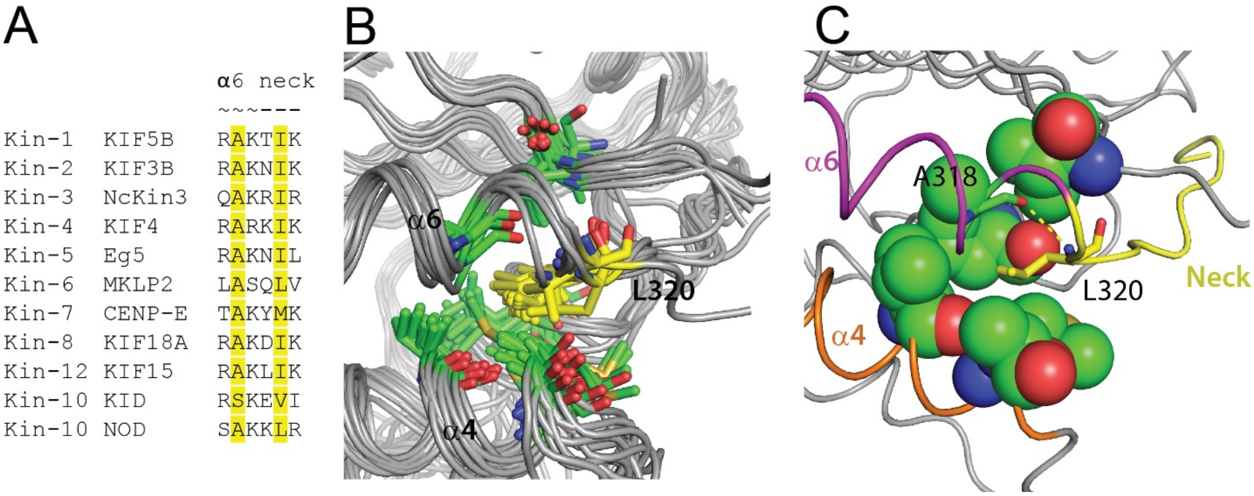


**Figure S7. Homology model of putative neck linker docking of NOD** A) Alignment of the last three residues of α3 and the first three residues of the neck of various kinesin subfamilies. The backbones of the highlighted residues form a hydrogen bond that orients the hydrophobic residue (L320 in NOD) to orient itself inside the hydrophobic pocket (V9,V260,M263,A264,L285,A317 in NOD). B) Fourteen different homology models of NOD were generated from template structures with docked neck linkers (see methods). In all cases the side chain of L320 (yellow) is oriented inside the hydrophobic pocket (green = carbon, red = oxygen, blue = nitrogen, gold = sulfur). C) Representative homology model where L320 forms a hydrogen bond with A318 in α6 which orients the side chain inside the hydrophobic pocket (yellow = neck linker, purple = α6, orange = α4, spheres = neck linker binding pocket where green = carbon, red = oxygen, blue = nitrogen, gold = sulfur.

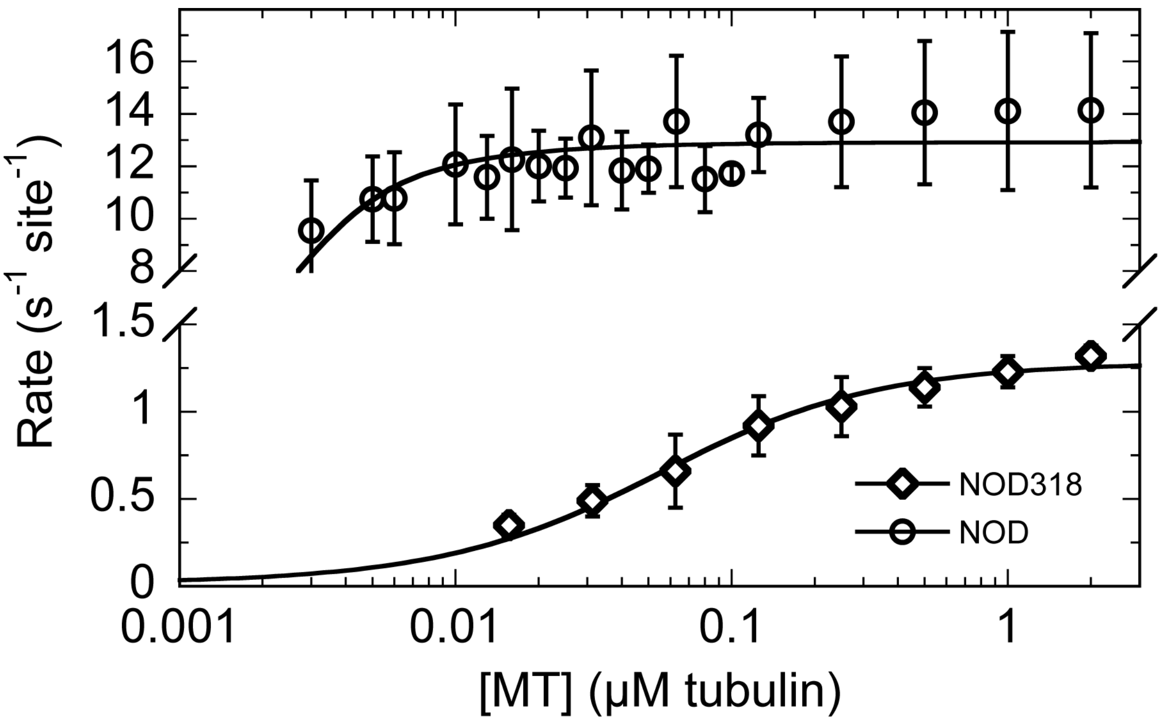


**Figure S8. Microtubule dependent steady-state ATPase of NOD318 compared to NOD.** Experiments with NOD318 were performed at low ionic strength. NOD = 0.02 µM. Error bars represent the standard deviation across independent experiments (n = 2). Reported error for each constant represents the standard error obtained from non-linear least squares analysis. NOD318 (◊): $k_{cat}$ = 1.27 ± 0.03 s$^{-1}$; $K_{0.5,\mathrm{MT}}$ = 0.045 ± 0.006 µM. NOD (○): $k_{cat}$ = 12.8 ± 0.3 s-1; $K_{0.5,\mathrm{MT}}$ = 0.0005 ± 0.0002 µM NOD data (see Figure 1C) is included as reference.

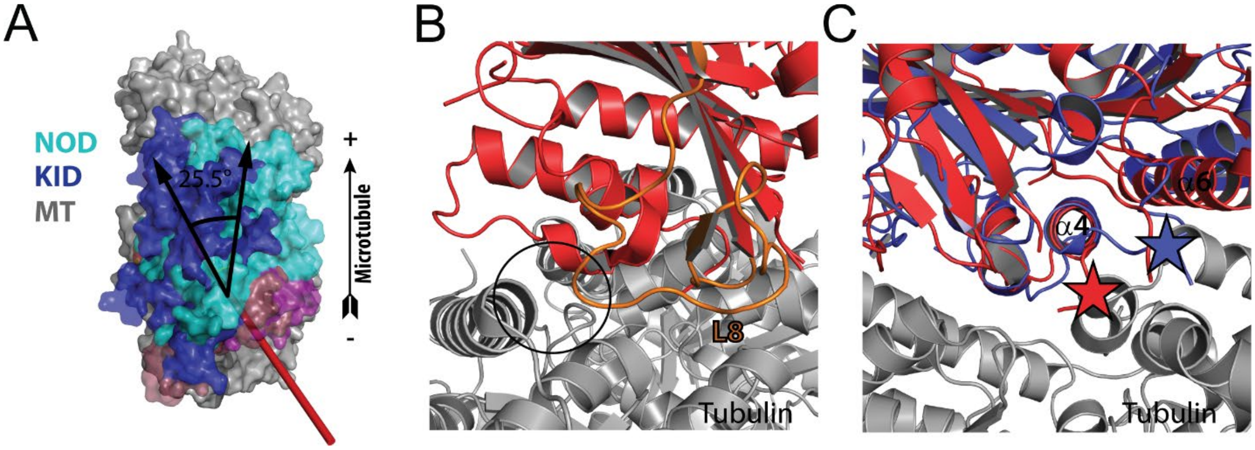


**Figure S9. NOD and KID docked onto the microtubule structure.** Structures were docked onto the microtubule lattice by aligning helix α4 of NOD and KID with kinesin-1 bound to the microtubule (3J8X). A) The N-terminal and upper subdomains of NOD are rotated 25.5° clockwise compared to KID. B) NOD is able to rotate while avoiding clashes with the microtubule lattice due to its missing hairpin in L8. C) In both structures (red = NOD; blue = KID) clashes between the lower subdomain and the microtubule lattice are observed.

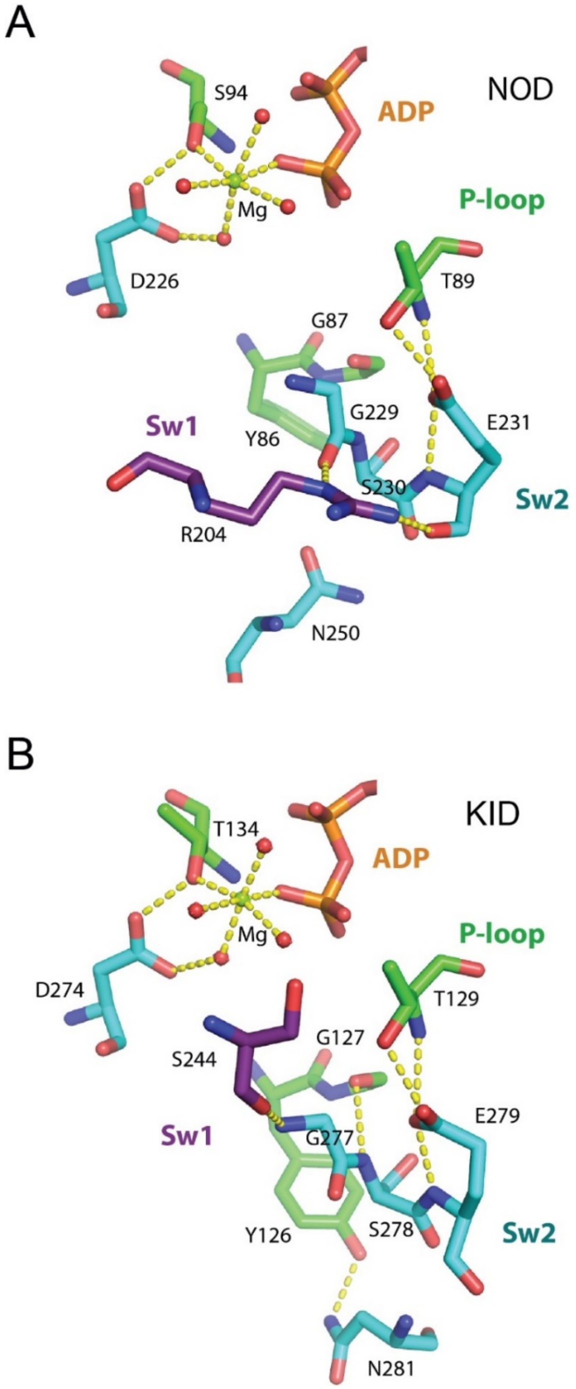


**Figure S10. Active site structure of NOD and KID.** Hydrogen bond configuration between switch-1, switch-2, and the P-loop in the active site of KID (A) and NOD (B). The switch-2 loop in both kinesins make two conserved interactions with the P-loop (NOD: D226-S94, E231-T89; KID: D274-T134, E279-T129) while KID has two additional interaction (S278-G127 and N281-Y126). The switch-1 and switch-2 of KID have one point of contact (G277-S244) which is conserved in ADP bound structures across the kinesin superfamily.
